# Integrated evaluation of telomerase activation and telomere maintenance across cancer cell lines

**DOI:** 10.1101/2021.01.22.426137

**Authors:** Kevin Hu, Mahmoud Ghandi, Franklin W. Huang

## Abstract

In cancer, telomere maintenance is critical for the development of replicative immortality. Using genome sequences from the Cancer Cell Line Encyclopedia and Genomics of Drug Sensitivity in Cancer Project, we calculated telomere content across 1,299 cancer cell lines. We find that telomerase reverse transcriptase (*TERT*) expression correlates with telomere content in lung, central nervous system, and leukemia cell lines. Using CRISPR/Cas9 screening data, we show that lower telomeric content is associated with dependency of CST telomere maintenance components. Increased dependencies of shelterin members are associated with wild-type *TP53* status. Investigating the epigenetic regulation of *TERT*, we find widespread allele-specific expression in promoter-wildtype contexts. *TERT* promoter-mutant cell lines exhibit hypomethylation at PRC2-repressed regions, suggesting a cooperative global epigenetic state in the reactivation of telomerase. By incorporating telomere content with genomic features across comprehensively characterized cell lines, we provide further insights into the role of telomere regulation in cancer immortality.

## Introduction

Telomeres, repetitive nucleoprotein complexes located at chromosomal ends, are an important component of genomic stability (De Lange, 2009). As protective chromosomal caps, telomeres prevent potentially lethal end-fusion events (McClintock, 1941, 1942) and mis-processing of chromosomal ends as damaged sites by the DNA repair machinery (De Lange, 2005; Verdun & Karlseder, 2007). Due to factors such as incomplete DNA replication and oxidative stress, telomeres gradually shorten with successive rounds of cell division (Olovnikov, 1973). If left unchecked, telomere attrition eventually triggers growth arrest and senescence, and further shortening can lead to acute chromosomal breakage and cell death. Telomere shortening therefore acts as a major obstacle in the course of tumor development (Hackett & Greider, 2002), and inhibition of telomere maintenance offers still largely untapped opportunities for targeted cancer therapies (Damm et al., 2001; Dikmen et al., 2005; Flynn et al., 2015).

Telomere shortening in embryonic development and in certain adult cell populations is offset by telomerase (Greider & Blackburn, 1985), a ribonucleoprotein enzyme with a core reverse transcriptase, *TERT*, that lengthens telomeres by catalyzing the addition of TTAGGG nucleotide repeats from an inbuilt RNA template component, *TERC* (Feng et al., 1995; Yu et al., 1990). Although telomerase is transcriptionally silenced in the majority of somatic cells, telomerase is reactivated in over 85% of all human cancers (N. W. Kim et al., 1994).

Reactivation of telomerase is associated with a diverse set of genomic alterations, the most common of which include highly recurrent mutations in the *TERT* promoter (Horn et al., 2013; Huang et al., 2013), aberrant methylation (D. D. Lee et al., 2019) or copy number amplification (Zhang et al., 2000) of *TERT*, and modulation of the numerous transcription factors that regulate *TERT* expression (Greider, 2012; Wu et al., 1999). Of the minority of cancers that do not reactivate telomerase, many depend upon alternative lengthening of telomeres (ALT), a process that exploits mechanisms of homologous recombination and is characterized by heterogeneous telomere lengths, mutations in the *ATRX* and *DAXX* chromatin-regulating factors, and genome instability (Cesare & Reddel, 2010).

The readily identifiable nature of telomeric DNA repeats has motivated the development of computational methods for the determination of telomere content from whole-genome sequencing (WGS) and whole-exome sequencing (WES) data (Ding et al., 2014). Recently, such methods were employed to characterize telomere content across tumor sequencing data from The Cancer Genome Atlas (TCGA) and the Genotype-Tissue Expression (GTEx) project, which identified genomic markers of relative telomere lengthening (Barthel et al., 2017; Castel et al., 2019). To gain a greater functional understanding of the landscape of telomere maintenance in cancer, we estimated telomeric DNA content (subsequently referred to as telomere content (Feuerbach et al., 2019)) across a diverse array of human cancer cell lines profiled in the Cancer Cell Line Encyclopedia (CCLE) (Barretina et al., 2012; Ghandi et al., 2019) and Genomics of Drug Sensitivity in Cancer (GDSC) (Yang et al., 2013) projects. We hypothesized that telomere content could be reflective of underlying mechanisms of attrition, maintenance, and repair, which may be reflected in associations with genetic markers. By combining these estimates with a rich set of existing CCLE annotations, we determined genetic, epigenetic, and functional markers of telomere content and telomerase activity across a diverse panel of human cancer cell lines.

## Results

### Telomere content across cancer cell lines

Telomeric reads can be identified in DNA-sequencing reads using the canonical tandemly repeated TTAGGG motif, and normalized telomeric read counts may provide an accurate estimate of telomere content (Ding et al., 2014). We quantified telomere content across cell lines using WGS and WES data from the CCLE (Barretina et al., 2012; Ghandi et al., 2019) and GDSC (Yang et al., 2013) datasets. In particular, we considered 329 cell lines profiled with WGS and 326 with WES in the CCLE, and 1,056 samples profiled with WES in the GDSC, of which 55 were non-cancerous matched-normal samples. We note that our estimates state telomeric DNA repeat tract content, which is a normalized measure of telomeric reads in a sample, rather than solely telomere length, because telomere length requires the identification of true telomeric DNA from intrachromosomal, non-terminal telomeric DNA repeat tracts and extrachromosomal telomeric DNA (Feuerbach et al., 2019). To assess the fidelity of our telomere content measurements, we examined the agreement between the telomere content estimates in overlapping cell lines from different sequencing datasets (**Supplementary Fig. 1**). We observed high agreement between the telomere content estimates derived from CCLE WGS and GDSC WES data (*r* = 0.84, *P* = 3.7×10^−79^, *n* = 286) and moderate agreement between CCLE WGS and CCLE WES estimates (*r* = 0.71, *P* = 1.5×10^−6^, *n* = 36). Therefore, we generated a merged telomere content dataset (**Supplementary Methods**) by combining the normalized log-transformed telomere contents derived from the CCLE WGS and GDSC WES datasets for downstream analyses.

The overall distribution of telomere content displayed a slight skew (**Supplementary Fig. 1**) towards longer telomeres, perhaps reflective of cell lines dependent upon ALT, a hallmark of which is telomeres of abnormal and heterogeneous lengths (Bryan et al., 1997; Heaphy, Subhawong, et al., 2011). We matched a substantial number of cell lines (282 for CCLE WGS, 554 for GDSC WES) with the age of the donor at the time of removal, from which we observed weak negative (vs. CCLE WGS: *r* = -0.05, *P* = 0.39; vs. GDSC WES: *r* = -0.17, *P* = 6.0×10^−5^) correlations between telomere content and the age of the original donor (**Supplementary Fig. 2a**). Among 1,099 merged CCLE WGS and GDSC WES samples, we found raw telomere content to vary substantially both between (*P* = 2.0×10^−15^, Kruskal-Wallis *H* test) and within (**Fig. 1**) cell lines of different primary sites. Cell lines of hematopoietic origin (namely leukemias and lymphomas, which comprised 156 lines) tended to have higher telomere contents on average (*P* = 2.0×10^−8^, two-sided Mann-Whitney *U* test), perhaps due to their elevated levels of telomerase expression, which were the highest among all subtypes. The 55 non-cancerous samples profiled as part of the GDSC also displayed relatively high telomere contents (*P* = 5.9×10^−4^, two-sided Mann-Whitney *U* test), consistent with previous reports of widespread telomere shortening in cancer (Barthel et al., 2017). The greatest average telomere content, however, was found across 44 bone/osteosarcoma cell lines (*P* = 7.2×10^−5^, two-sided Mann-Whitney *U* test; **Fig. 1**), likely because the rate of ALT is relatively high in bone cancers (Scheel et al., 2001). The cell line with the highest telomere content was the U2-OS osteosarcoma line, a well-characterized model for ALT (Bryan et al., 1997).

**Figure 1.**
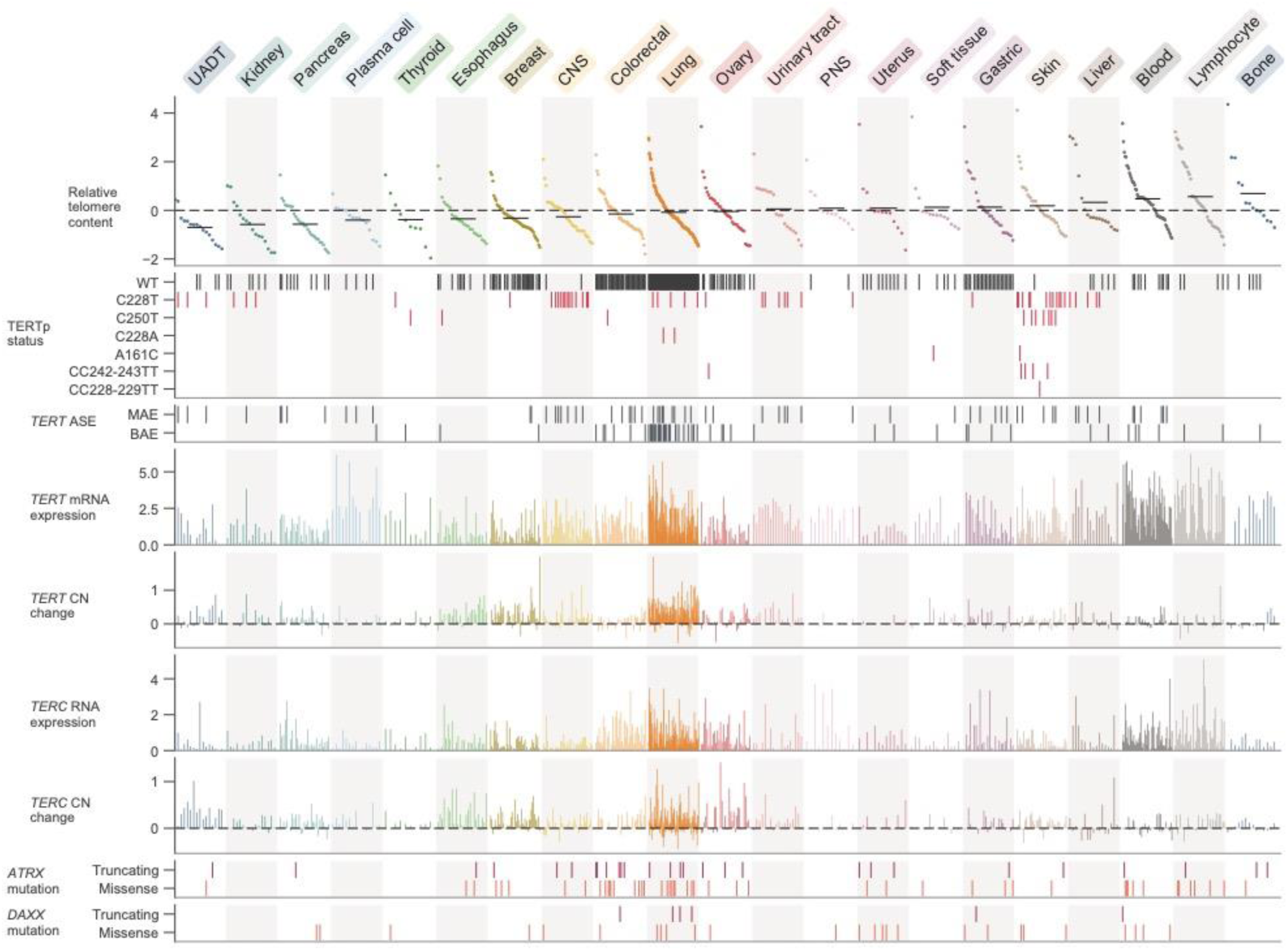
Telomere content and related genomic features across human cell lines. Cell lines were grouped by cancer type and ordered by telomere content within each type, and are displayed such that each column represents a cell line. Telomere content measurements reflect combined *z*-scored estimates derived from CCLE WGS and GDSC WES with means for samples with telomere content estimates from both sources. Relative copy number values are shown as log_2_(relative to ploidy + 1) - 1. Cell lines shown are filtered such that annotations for telomere content, *TERT* and *TERC* RNA-seq expression, *TERT* and *TERC* copy number, and *ATRX* and *DAXX* mutation status are all available, and each cancer type is represented by at least ten cell lines (*n* = 683 cell lines total). RNA expression estimates are in terms of log_2_(TPM+1). CNS: central nervous system; PNS, peripheral nervous system; UADT, upper aerodigestive tract.

### Genomic alterations associated with telomere content

To characterize the genomic signatures of telomere content, we correlated the CCLE WGS and GDSC WES telomere content estimates against molecular annotations in the CCLE, with a focus on alterations known to be associated with telomere maintenance. First, we observed a positive, though weak, correlation between telomere content and *TERT* mRNA levels as determined by RNA sequencing (RNA-seq) both within each subtype (**Supplementary Fig. 4a,b**) and overall (**Supplementary Fig. 4c,d**), although this was not an outlier compared against the expression levels of all genes (**Supplementary Fig. 3b**). In specific cancer types such as central nervous system (CNS), lung, and leukemia, we found a higher correlation between *TERT* mRNA expression and telomere content (**Supplementary Fig. 4a,b**). Furthermore, we found negative associations between *TERT* mRNA levels and telomere content in bone and peripheral nervous system cell lines (**Supplementary Fig. 4a,b**). Because ALT is most commonly found in these cancer types, this may be a consequence of the near-mutual exclusivity between *TERT* expression and markers of ALT (Killela et al., 2013; M. Lee et al., 2018). Although mutations in *ATRX* and *DAXX* are closely associated with the development of the ALT phenotype (Brosnan-Cashman et al., 2018; Clynes et al., 2015; Heaphy, De Wilde, et al., 2011; Lovejoy et al., 2012; Ramamoorthy & Smith, 2015), comparisons between telomere content and mutations in *ATRX* and *DAXX* yielded significant associations only between *DAXX* alterations and GDSC WES telomere content (**Supplementary Fig. 3a**). We further repeated association tests with *TP53, VHL*, and *IDH1* as identified previously among TCGA samples (Barthel et al., 2017), with which we confirm that truncating *VHL* mutations are associated with reduced telomere lengths (**Supplementary Fig. 3a**) although this may be confounded by the high occurrence of *VHL* mutations in kidney cell lines. Whereas we found relatively few significant associations between telomere content and mutations, we note that we were limited to an absolute estimate of telomere content as opposed to a relative measure of somatic telomere lengthening, which requires a paired normal sample (Barthel et al., 2017).

### Telomere content associates with CST complex dependencies

Having explored the association between telomere content and non-perturbative annotations, we next considered whether variations in telomere content could confer or reduce selective vulnerabilities to inactivation of certain genes. In particular, we hypothesized that telomere content may be associated with vulnerabilities to reductions in the levels of telomere-regulating proteins. To reveal such associations, we correlated our telomere content estimates with gene inactivation sensitivities assessed via genome-wide CRISPR-Cas9 (Avana (Meyers et al., 2017)) and RNAi viability screens (Achilles RNAi (Tsherniak et al., 2017) and DRIVE (McDonald et al., 2017)). Although we found no dependencies that displayed outlier associations with telomere content in the Achilles RNAi screen (**Supplementary Fig. 3b**), we discovered that sensitivity to Avana CRISPR-Cas9 knockouts of each of the three CST complex proteins was an outlier association with telomere content estimates computed from both the GDSC WES and CCLE WGS data (**Fig. 2a,b, Supplementary Fig. 5b**). Namely, increased sensitivities to knockout of the CST complex components, which are key mediators of telomere capping and elongation termination (Chen et al., 2012), were correlated with lower telomere content (**Supplementary Fig. 5a**). Although the CST complex was not assessed in the DRIVE screening dataset, we also found *TERF1*, a key shelterin component, to be among the positive correlated genes with telomere content in the DRIVE panel (**Supplementary Table 2**).

**Figure 2.**
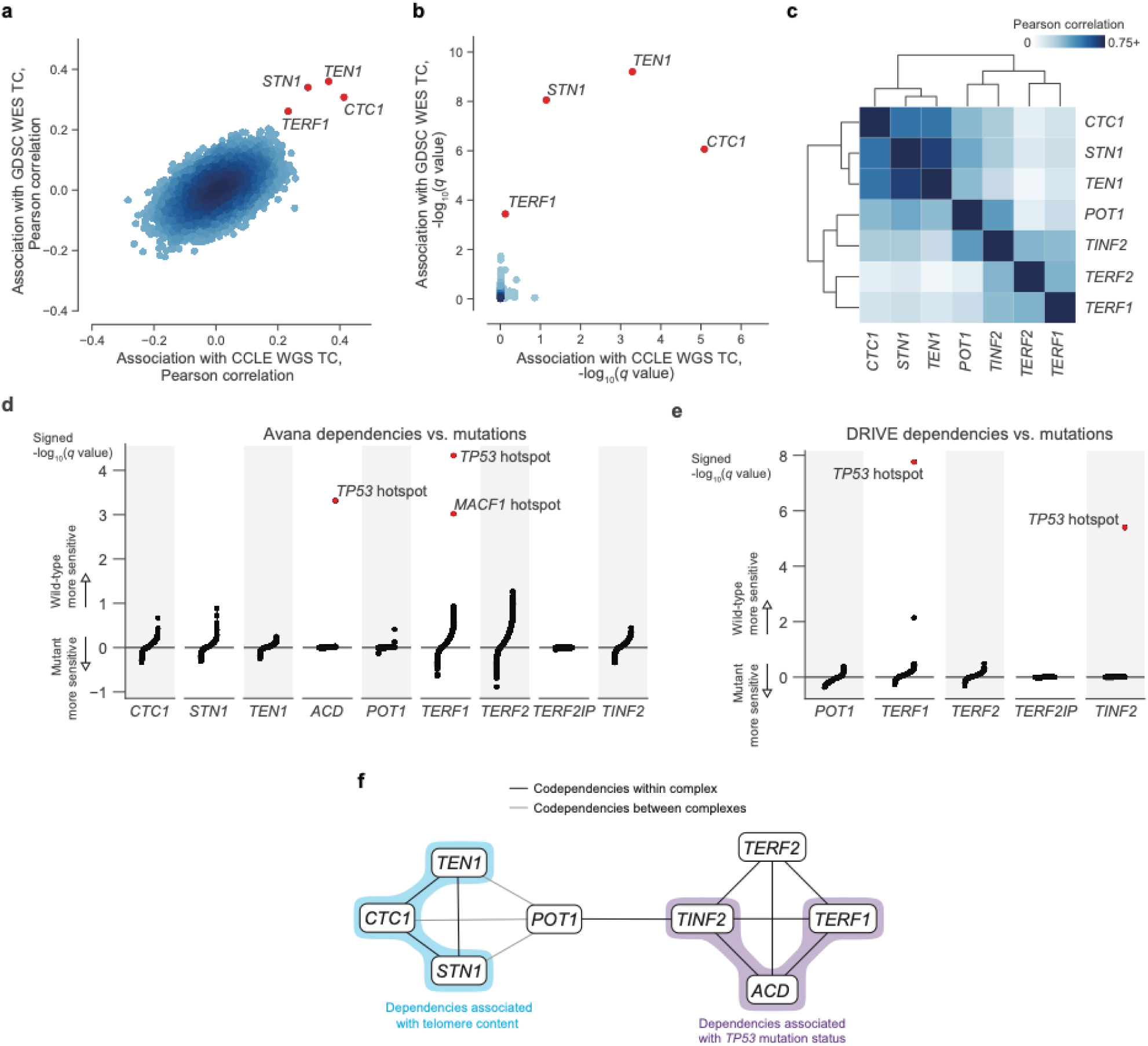
Telomere-binding protein dependencies are associated with telomere content and *TP53* mutation status. **a**, Pairwise plot of Pearson correlations between dependencies of all genes in the Avana dataset and CCLE WGS telomere content (*x*-axis, *n* = 192–210 cell lines) and GDSC WES telomere content (*y*-axis, *n* = 395–416 cell lines) estimates. **b**, Pairwise plot of significance levels of correlations shown in (**a**) with correction for multiple hypothesis testing. **c**, Pairwise Pearson correlation matrix between Avana dependencies among CST members and five shelterin components (*n* = 757-769 cell lines; **Supplementary Table 3**). **d**, Associations of CST and shelterin member Avana dependency scores with damaging and hotspot mutations (*n* = 756–767 cell lines). For each gene dependency, mutation associations are shown ranked by signed *q*-value, with sign indicating the direction of association (negative for greater sensitivity in mutants, positive for greater sensitivity in wild-type cell lines). *P* values determined using two-sided Pearson’s correlation test. **e**, Associations of shelterin member DRIVE dependency scores with damaging and hotspot mutations (*n* = 372–375 cell lines; **Supplementary Table 3**) under the same scheme used in (**d**). **f**, Network schematic of the co-dependency matrix shown in (**c**) and annotated with association with telomere content or *TP53* mutation status.

Using the CST complex as a seed set, we subsequently queried all dependencies under the premise that associated gene dependencies reflect coordinated functions (Pan et al., 2018). Within the Avana panel (*n* = 757–769), we found significant (FDR < 0.01) outlier associations between the CST complex genes and genes encoding five additional telomere-associating proteins (*ACD, POT1, TERF1, TERF2, TINF2*). These five additional proteins, together with *TERF2IP*, comprise the shelterin complex, the protector and regulator of telomere length and topology (De Lange, 2005). Interestingly, whereas the five other dependencies were positively associated with telomere content, *TERF2IP* displayed a weak negative association (**Supplementary Fig. 5c**), suggesting that *TERF2IP* may play a distinct regulatory role in shelterin function compared to the other members. To examine the dependency landscape of the CST complex and these five other telomere-associated proteins, we computed a correlation matrix involving these eight genes, clustering of which yielded two main subgroups: one comprised of the CST complex members, and another of the five other genes (**Fig. 2c**). Despite this separation, *POT1* and *TINF2* also displayed notable correlations with CST dependencies, possibly serving as the primary mediators of previously-reported functional interactions between the shelterin and CST complexes (Chen et al., 2012; Wan et al., 2009).

Whereas we found strong codependency relationships within this group of eight telomere-associated proteins, we also noticed that certain shelterin members displayed notable codependencies with p53 pathway members such as *MDM2, ATM*, and *TP53* itself (**Supplementary Fig. 6a,b**). Because sensitivity to perturbation of the p53 pathway is highly associated with *TP53* mutations in cancer (McDonald et al., 2017), we asked if these codependency relationships were also associated with hotspot mutations in *TP53*. In fact, *TP53* was a significant (*FDR* < 0.001) outlier when a comprehensive set of hotspot and damaging mutations was compared against sensitivity to *ACD* and *TERF1* dependencies in the Avana panel (**Fig. 2d, Supplementary Fig. 6c**) and against *TERF1* and *TINF2* dependencies in the DRIVE panel (**Fig. 2e, Supplementary Fig. 6d**). These links between these gene dependencies and *TP53* mutation status reprise and extend previous reports of p53-dependent DNA damage responses to *TERF1* and *TINF2* depletion (Pereboeva et al., 2016; Rosenfeld et al., 2009). Taken together, we find that CST and shelterin dependencies are correlated with each other, telomere content, and *TP53* mutation status (**Fig. 2f**).

### Investigation of *ATRX*-*DAXX* dependencies

To further explore the landscape of telomere regulation from a dependency standpoint, we also asked if there were dependency associations surrounding the ATRX-DAXX histone chaperone complex, which is often depleted in ALT tumors (Brosnan-Cashman et al., 2018; Clynes et al., 2015; Heaphy, De Wilde, et al., 2011; Lovejoy et al., 2012). *ATRX* and *DAXX* dependencies were found to be closely associated in the Avana dataset (**Fig. 3a**), with both genes being the top codependency with the other. Although these dependencies were not present as top correlates against telomere content, we reasoned that they may be associated with other genomic markers. In fact, among all genes, *PML*, which has previously been implicated in the organization and regulation of ATRX and DAXX, was the top gene expression correlate with *DAXX* dependency (**Fig. 3b**). To further investigate the genomic markers of these dependencies, we compared *ATRX* and *DAXX* dependencies to CCLE quantitative proteomics measurements (Nusinow et al., 2020). We found an outlier association between reduced *DAXX* dependency and increased protein levels of ZMYM3 (**Fig. 3c**), a chromatin-binding histone deacetylase complex member involved in the DNA damage response (Leung et al., 2017). Although *ATRX* and *ZMYM3* are located on adjacent bands of the X chromosome, ATRX protein levels were not significantly associated with *DAXX* dependencies. Interestingly, a complementary association of ZMYM3 protein levels and all Avana gene dependencies not only revealed *DAXX* as the top associate, but also a significant association with *TERF1* dependency (**Fig. 3d**). Taken together, these results suggest additional modulators of the ATRX-DAXX axis and connections between *DAXX, ZMYM3*, and *TERF1* regulation.

**Figure 3.**
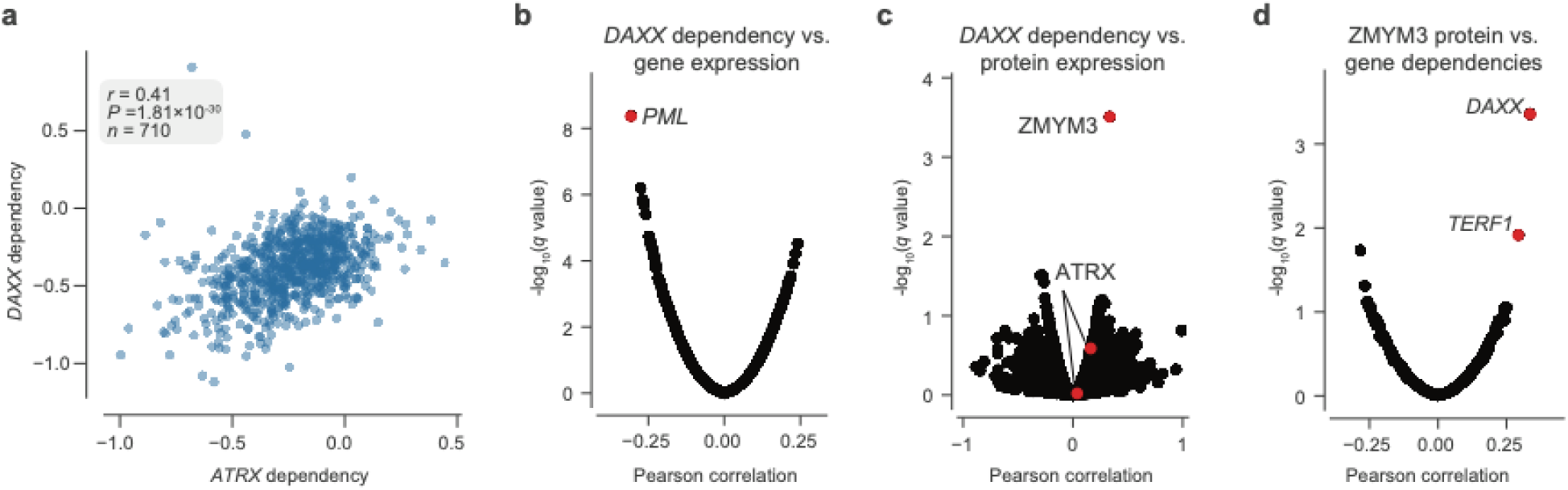
Transcriptomic and proteomic determinants of *DAXX* dependency. **a**, Comparison of *DAXX* and *ATRX* dependency levels as measured in the Avana dataset. **b**, Correlations of *DAXX* Avana dependencies with genome-wide RNAseq gene expression estimates (*n* = 566 cell lines). **c**, Correlations of *DAXX* Avana dependencies with all profiled protein expression estimates (*n* = 4–262 cell lines). Two ATRX isoforms (UniProtKB: P46100 and P46100-6) are also highlighted. **d**, Correlations of ZMYM3 protein levels with all Avana dependencies (*n* = 242–263 cell lines). *P* values determined using two-sided Pearson’s correlation test.

### Patterns and mechanisms of telomerase expression

Having thoroughly characterized telomere content and its related dependencies across the CCLE, we next focused on the regulation of *TERT* transcription. Across 1,019 samples previously profiled with deep RNAseq, we found that hematopoietic cell lines (leukemias, lymphomas, and myelomas) were associated with the greatest mean expression of *TERT* (*P* = 5.8×10^−26^, two-sided Mann-Whitney *U* test; **Fig. 1**). In contrast, *TERT* expression was significantly reduced (*P* = 2.0×10^−22^, two-sided Mann-Whitney *U* test) and generally undetectable in non-cancerous fibroblast cell lines. With regards to the telomerase RNA component (*TERC*), we found strong associations between *TERC* RNA expression and RNA levels of several small Cajal body-specific RNAs (scaRNAs) and histone subunit RNAs (**Supplementary Fig. 4e**). The co-expression of *TERC* and these other RNAs may be a consequence of their shared localization, processing, and regulation in Cajal bodies (Gall, 2003; Venteicher et al., 2009; Zhu et al., 2004), as *TERC* itself contains an H/ACA box small nucleolar RNA (snoRNA) domain (Mitchell et al., 1999) and is a scaRNA family member. Moreover, these associations suggest that variations in Cajal body processing may act as factors in *TERC* reactivation (Cao et al., 2008). Although we focused on the transcriptional features of telomerase, it is important to note that other factors in addition to *TERT* and *TERC* expression determine telomerase activity and the eventual maintenance of telomere length (Listerman et al., 2013). Despite these orthogonal factors, *TERT* enzymatic activity remains strongly correlated with raw levels of *TERT* expression (Takakura et al., 1998).

Widespread transcriptional reactivation of *TERT* in cancer is driven by a variety of factors. Aside from copy number amplifications (Zhang et al., 2000), highly recurrent mutations in the *TERT* promoter drive strong monoallelic *TERT* re-expression (Bell et al., 2015; Huang et al., 2015). Using WGS and targeted sequencing of the *TERT* promoter provided in the CCLE, we considered *TERT* promoter status for 503 cell lines. We found that only the C228T (chr5:1,295,228 C>T) mutation was significantly (*P* = 2.8×10^−5^, two-sided Mann-Whitney *U* test) associated with an increase in TERT expression (**Supplementary Fig. 8b**). Surprisingly, the mean level of *TERT* expression in monoallelic contexts was only slightly lower than that of biallelic contexts, with less than a 1.5-fold difference between the groups (*P* = 0.03, two-sided Mann-Whitney *U* test). Given that cells with biallelic *TERT* expression do so with twice the transcriptional source sites as those with monoallelic *TERT* expression, this reduced difference may be a consequence of the effects of the *TERT* promoter mutation in producing particularly robust monoallelic expression (Huang et al., 2013) or expression of TERT from multiple sites all of the same allele (Rowland et al., 2019).

To further explore allele-specific expression (ASE) patterns of *TERT*, we employed an ASE-calling pipeline (**Supplementary Methods**) and determined *TERT* allele-specific expression status for 157 cell lines (**Supplementary Fig. 7a**), an increase of 69 cell lines compared to a previous report using CCLE WGS data (Huang et al., 2015). Out of these 157 cell lines, 87 express *TERT* from a single allele. Moreover, of these 157 cell lines, 129 have a sequenced promoter, with which we confirm that promoter mutations unanimously drive monoallelic expression (**Supplementary Fig. 7c**). Our expanded set of cell lines also reveals several new tissues of origin in which *TERT* is monoallelically expressed without a mutant promoter, such as hematopoietic cell lines (**Supplementary Fig. 7a,b**). This high proportion of *TERT* monoallelic expression then led us to ask whether there are genomic alterations aside from promoter mutations that could lead to ASE. Under the assumption that such alterations may also induce ASE in a larger region than a promoter mutation, we determined ASE status in *SLC6A19* and *CLPTM1L*, which are the most immediate neighbor genes of *TERT*. Because the number of samples with annotated ASE in both *TERT* and these neighbors was not large enough for comparisons of overlapping ASE, we instead examined the individual frequencies of ASE among these three genes. Compared to promoter-wildtype monoallelic *TERT* expression occurring in 36% (47 of 129) of samples, only 9.4% (11 out of 117) of samples expressed *CLPTM1L* from a single allele and 22% (5 of 22) expressed *SLC6A19* from a single allele, and we were unable to assess co-occurrence due to lack of overlap in samples with heterozygous SNPs in both *TERT* and *CLPTM1L* or *SLC6A19*. Furthermore, a search for structural variants in the surrounding 100 kilobase regions yielded no significant associations. However, given that only 106 cell lines had both a known *TERT* ASE status and the required WGS data for structural variant determination, more genomic annotations may be needed for the discovery of additional mechanisms driving the monoallelic expression of *TERT*.

### Distinct TERTp Methylation Patterns at the *TERT* locus

Aside from monoallelic expression, *TERT* promoter mutations are characterized by unique patterns of epigenetic marks, namely allele-specific CpG methylation (ASM) and H3K27me3 repressive histone modifications (Stern et al., 2015, 2017) and long-range chromatin interactions (Akincilar et al., 2016). Using genome-wide RRBS data across 928 cell lines, we elucidated associations between CpG-site methylation of the *TERT* locus (namely, *TERT* and the surrounding 5 kilobases) and *TERT* promoter mutations. Examination of methylation patterns at the *TERT* locus revealed five prominent ASM clusters in the *TERT* locus, corresponding roughly to the upstream 5kb region (containing the promoter), part of a CpG island overlapping the first exon, two other parts of this CpG island, and the remaining gene body (**Fig. 4a**). Comparison of each region’s mean ASM against TERTp mutant status revealed that TERTp mutants exhibited strong and significant (*P* < 0.01) increases in ASM in the promoter (*n* = 485), remaining gene body (*n* = 493), and exon 1 (*n* = 478) regions (**Fig. 4b, Supplementary Fig. 8a,b**). In contrast, TERTp wild-type cell lines tended to lack ASM throughout the *TERT* locus, instead being hypermethylated in all regions except for exon 1 (**Fig. 4b**), and partial hypomethylation in promoter mutants may reflect the hemizygous methylation previously observed at the *TERT* locus (Rowland et al., 2020; Stern et al., 2015, 2017). Absolute methylation of the remaining gene body was positively correlated with *TERT* expression in both promoter status contexts (**Supplementary Fig. 8d**), which parallels previous reports of a positive correlation between *TERT* expression and methylation (Barthel et al., 2017; Salgado et al., 2019). Exon 1 methylation was elevated in nearly all cell lines with monoallelic *TERT* expression in both the mutant promoter context (*P* = 6.3×10^−3^, two-sided Mann-Whitney *U* test, *n* = 81) and the wildtype one (*P* = 1.6×10^−4^, two-sided Mann-Whitney *U* test, *n* = 98) compared to biallelic *TERT* expressors (Stern et al., 2017). Interestingly, although most cell lines with monoallelic *TERT* expression displayed partially elevated methylation levels in exon 1 (**Fig. 4b**), only promoter-mutant cell lines were hypomethylated in the surrounding regions, suggesting that the epigenetic state of promoter mutants is in fact distinct from that of promoter-wildtype monoallelic *TERT* expressors.

**Figure 4.**
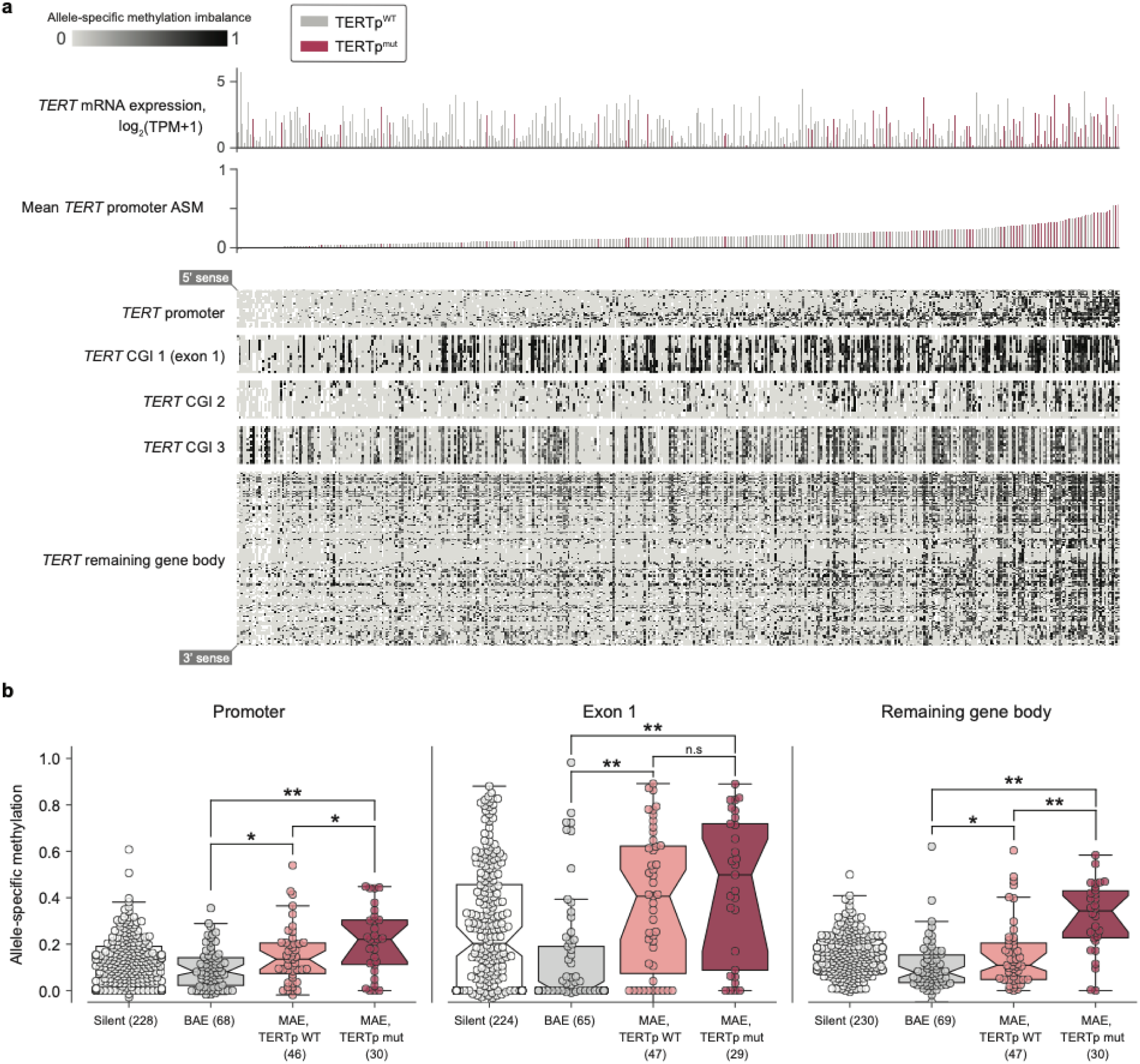
Allele-specific methylation of the *TERT* locus is indicative of both promoter mutation status and allele-specific expression. **a**, Heatmap of CpG methylation levels along the *TERT* locus, sorted in order of mean methylation levels along the upstream 5kb region. *TERT* gene expression levels are also indicated for each cell line. Each column represents a cell line (*n* = 451), and each row represents a CpG pair (*n* = 208) sorted from the 5’ to 3’ direction along the *TERT* sense strand. White blocks indicate missing ASM values. Cell lines with unavailable ASM values for at least half of *TERT* locus CpGs were excluded. **b**, ASM levels of *TERT* locus subregions in cell lines are indicative of TERTp status and allele-specific expression. Boxes, interquartile range (IQR); center lines, median; whiskers, maximum and minimum or 1.5 × IQR; notches, 95% confidence interval of bootstrapped median using 1,000 samples and a Gaussian-based asymptotic approximation. ^*^*P* < 0.05, ^**^*P* < 0.01, n.s, not significant; two-sided Mann-Whitney *U* test.

### Genome-wide epigenetic patterns in TERTp mutants

The observation that *TERT* promoter mutants display a hypomethylated *TERT* locus even compared to other monoallelic *TERT* expressors led us to ask if additional epigenetic signals are indicative of *TERT* promoter status. In particular, we considered the possibility that epigenetic changes to the *TERT* locus could in fact act as a cooperative factor (W. Kim et al., 2016; W. Kim & Shay, 2018) in tumorigenesis or tumor cell maintenance rather than as a consequence of *TERT* promoter mutations. To address this hypothesis, we performed a genome-wide search for CpG islands (CGIs) with significant differences in methylation levels in *TERT*p mutant cell lines compared to *TERT*p wild-type ones. If *TERT* hypomethylation were a downstream consequence of *TERT* promoter mutations, then we would expect *TERT* hypomethylation to be an isolated event, and thus there would be few CGIs outside the vicinity of *TERT* with methylation levels correlated with TERTp mutant status. Surprisingly, we instead found a broad genome-wide distribution of CGIs that were hypomethylated in TERTp^mut^ samples relative to TERTp^WT^ samples (**Fig. 5a**). Moreover, when correlated with a panel of global histone modification levels, we found that *TERT*p mutants exhibited increased levels of H3K9ac1K14ac0 and H3K9ac1K14ac1 marks (**Fig. 5c**), which have been suggested as marks of transcriptionally active chromatin (Ruthenburg et al., 2007). Likewise, when H3K9ac1K14ac0 levels were compared against a genome-wide panel of CGI ASM levels, the *TERT* CGI (chr5:1,289,275-1,295,970) was the top correlate (**Fig. 5d**).

**Figure 5.**
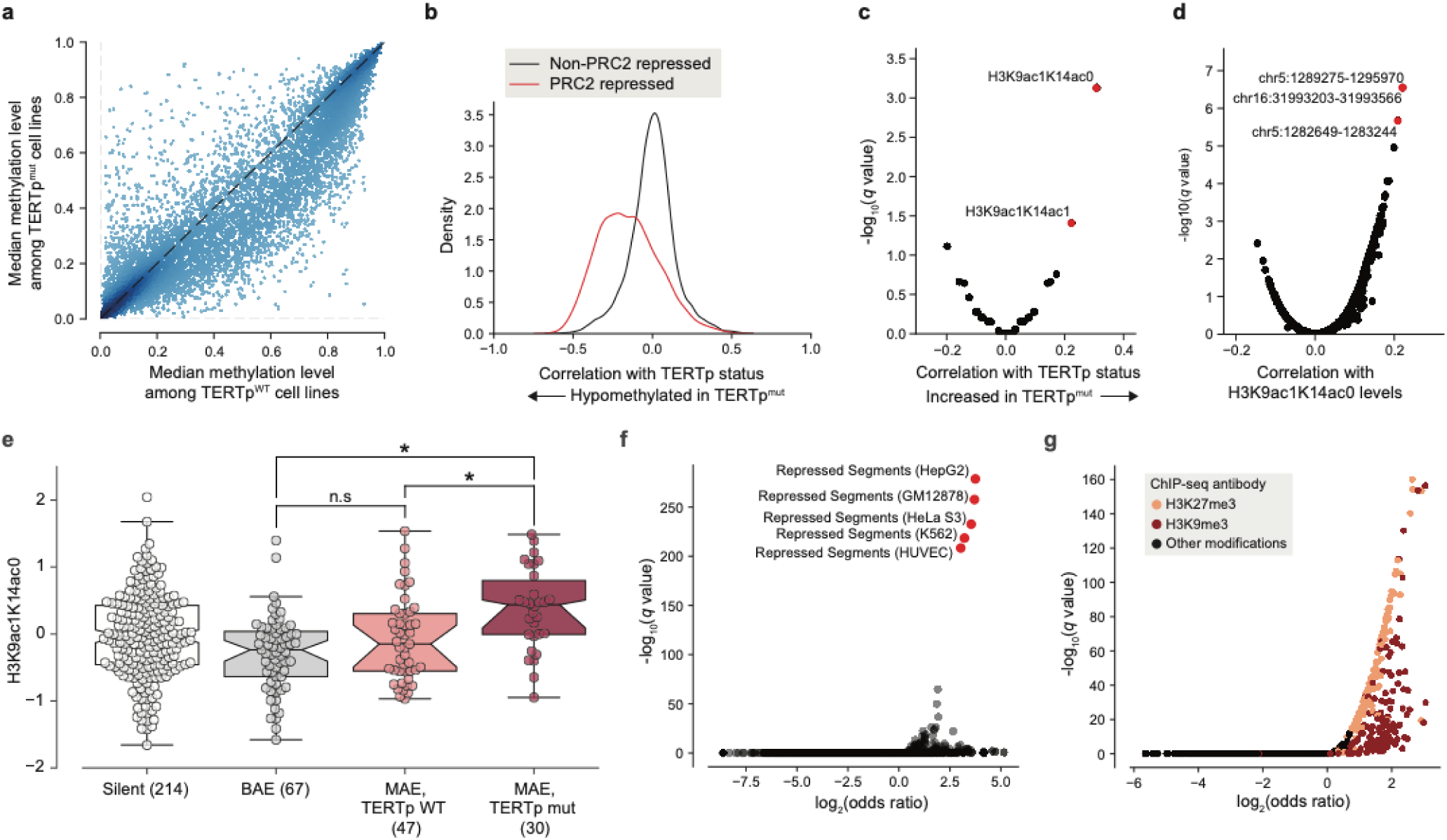
*TERT* promoter mutations associate with genome-wide decreased methylation of PRC2-repressed regions. **a**, Pairwise plot of median CGI methylation levels in TERTp^mut^ cell lines (*n* = 21–83; **Supplementary Table 6**) versus TERTp^WT^ cell lines (*n* = 95–410, **Supplementary Table 6**). **b**, Kernel density distributions of rank-biserial correlations between CGI methylation levels for PRC2-overlapping regions and non-PRC2-overlapping regions. A negative correlation indicates that a CGI is hypomethylated in TERTp^mut^ cell lines relative to TERTp^WT^ ones, and a positive correlation indicates the opposite. PRC2 regions were sourced from the HepG2 segmentation. **c**, Rank-biserial correlations between TERTp status (mutant or wild-type) and global histone modification levels (*n* = 302–475). Significance determined by two-sided Mann-Whitney *U* test. **d**, Pearson correlation levels between global H3K9ac1K14ac0 levels and ASM imbalance of CGIs (*n* = 261–884). **e**, H3K9ac1K14ac0 levels are significantly increased in TERTp mutants. Boxes, interquartile range (IQR); center lines, median; whiskers, maximum and minimum or 1.5 × IQR; notches, 95% confidence interval of bootstrapped median using 1,000 samples and a Gaussian-based asymptotic approximation. ^*^*P* < 0.01, n.s, not significant; two-sided Mann-Whitney *U* test. **f**, LOLA core set enrichment analysis of CGIs hypomethylated in TERTp^mut^ cell lines reveals enrichment of PRC2-repressed regions. **g**, LOLA ENCODE Roadmap region enrichment analysis of CGIs hypomethylated in TERTp^mut^ cell lines reveals enrichment of H3K9me3 and H3K27me3 regions.

To better understand the distribution of these TERTp^mut^-hypomethylated CGIs, we utilized Locus Overlap Analysis (LOLA) (Nagraj et al., 2018; Sheffield & Bock, 2016) to query the significance of overlaps between these CGIs and predetermined region sets. Among the top 1,000 TERTp^mut^-hypomethylated CGIs (**Fig. 5a**), we found significant (*FDR* < 0.0001) and robust 10-fold enrichment for polycomb repressive complex 2 (PRC2)-repressed regions (**Fig. 5f**) previously characterized in several cell lines (HepG2, GM12878, HeLa-S3, K562, and HUVEC). Beyond these top 1,000 hypomethylated CGIs, CGIs overlapping with PRC2-repressed segments were broadly hypomethylated in TERTp^mut^ cell lines and accounted for nearly all of the previously observed skew towards hypomethylation (**Fig. 5b**). Interestingly, the enrichment of PRC2 segments was much smaller (around 3.5-fold) in the remaining profiled cell line, H1-hESC. Against ENCODE ChIP-seq peak region sets, we also found significant overlap with the H3K9me3 and H3K27me3 heterochromatin marks (**Fig. 5g**). Furthermore, we also observed a moderate 2-fold (*P* = 5.1×10^−24^, Fisher’s exact test) enrichment for regions within ten megabases of most telomeres, consistent with previous reports that PRC2-repressed and H3K27me3-marked regions are enriched in telomeric and subtelomeric regions (Rosenfeld et al., 2009). The enrichment of these hypomethylated regions among telomere-proximal regions may also be indicative of a recently-reported telomere position effect, which has been shown to affect the chromatin accessibility of the *TERT* locus (W. Kim et al., 2016) located close to the chromosome 5p telomere. Given that PRC2 has previously been shown to exhibit allele-specific binding to the methylated silent allele in TERTp^mut^ cell lines (Stern et al., 2017), this genome-wide pattern of hypomethylation at PRC2 sites suggests that background epigenetic events may interact with promoter mutations in facilitating *TERT* expression.

We then asked whether the reduced methylation levels at telomere-proximal sites of PRC2 repression in the TERTp^mut^ CCLE samples displayed a similar pattern across TCGA primary tumor samples. To test this hypothesis, we estimated CGI methylation levels across 878 TERT-expressing TCGA samples characterized with both the Illumina 450k array and with a previously determined *TERT* promoter status (Barthel et al., 2017). Consistent with previous analysis of *TERT* methylation levels at the cg11625005 methylation probe, we find that TERTp^mut^ samples tended to exhibit hypomethylation (**Supplementary Fig. 9a,b**). Among 13,547 CGIs, we again found an enrichment of hypomethylation of PRC2-overlapping CGIs (**Supplementary Fig. 9c**), although this was less prominent than previously noted in the CCLE. LOLA enrichment analysis for TERTp^mut^-hypomethylated CGIs in the TCGA likewise confirmed significant enrichments (*FDR* < 0.0001) of PRC2-repressed regions and associated histone modifications as the top enriched region sets (**Supplementary Fig. 9d,e**). However, the fold-enrichment was less (about 5-fold) than that observed in the CCLE and did not display any significant enrichment in telomere-proximal regions (*P* = 0.70, Fisher’s exact test). Although the skew towards hypomethylation in *TERT* promoter mutants among these TCGA samples was weaker than in the CCLE samples, this may be the result of the more heterogeneous nature of these primary TCGA samples as well as the differences in coverage between the Illumina 450k array and RRBS.

## Discussion

To investigate the nature of telomeres and their maintenance mechanisms in cancer, we applied a functional genomics approach towards understanding molecular relationships across cancer cell lines. We estimated relative telomere content across over a thousand cancer cell lines and thus provide a useful reference for further studies on cancer cell line characteristics that have not to date considered this feature. We show that cell line telomere content indeed varies with factors such as tissue type, *TERT* mRNA expression, and mutations in genes such as *DAXX* and *VHL*. Moreover, we discovered novel relationships between telomere content and dependencies of CST and shelterin complex members, which was enabled by the high overlap between the cell lines profiled by our estimates and by several loss-of-function screens (McDonald et al., 2017; Meyers et al., 2017; Tsherniak et al., 2017).

When correlated with genome-wide gene dependency estimates, we found that increased sensitivity (Meyers et al., 2017) to depletion of CST complex members correlates with shorter telomeres, likely a consequence of the critical roles of the CST complex in both telomere protection and in terminating telomere elongation (Chen et al., 2012). Our findings raise the possibility that targeting the CST complex may preferentially affect cancer cells that harbor shorter telomeres, and telomere content may be used as a biomarker of response in tumors.

Likewise, CST complex dependencies were positively associated with the dependencies of several shelterin complex components, reflecting their functional interactions. Among these shelterin complex members, we further find that the responses to their depletion are highly dependent upon the presence of a wild-type *TP53* gene, with *TP53* mutants displaying reduced sensitivity to depletion of *ACD, TERF1*, and *TINF2*. Additional studies are required to validate these associations and to assess why only certain members of the shelterin complex show this *TP53*-dependent sensitivity effect.

In addition to telomere content, we also investigated the landscape of telomere maintenance mechanisms, namely mechanisms of *TERT* reactivation, across cancer cell lines. The enrichment of *TERT* promoter mutations in certain tissues has inspired several explanations, and our findings in both the CCLE and TCGA suggest a specific epigenetic signature that may underlie this unique pathway of telomere maintenance. We found that in *TERT* promoter mutants, CpG islands were preferentially hypomethylated in PRC2-repressed regions located near telomeres, which may relate to previous reports of a long-range telomere position effect (W. Kim et al., 2016; Yuan et al., 2019) and of *TERT* expression necessitating specific chromatin states in promoter-wildtype and mutant samples (Salgado et al., 2019). Considering that normal tissues typically exhibit particularly low methylation of the *TERT* promoter (Salgado et al., 2019; Stern et al., 2017) and that PRC2 occupies the inactive allele in *TERT* promoter mutants (Stern et al., 2017), our genome-wide signature may relate to the latter part of the two-step mechanism proposed for TERTp mutation-driven telomerase upregulation (Chiba et al., 2017). Moreover, epigenetic mechanisms have been shown to produce synergistic effects with driver mutations in tumor evolution (Tao et al., 2019). Besides reflecting a direct cooperation with *TERT* expression, this signature raises the possibility that the “memory” of short telomeres may be preserved through these telomere-proximal hypomethylated regions. It may also be indicative of the stemness of cell lines, which has been proposed as a major factor in the proliferative advantage of *TERT* promoter mutations (Chiba et al., 2015). Future studies will be necessary to elucidate the nature of this epigenomic signature, how it impacts the regulation of telomerase expression, and the complexities of *TERT* expression beyond binary measures of allele-specificity (Rowland et al., 2019). Furthermore, incorporation of telomere content into studies using cancer cell lines may help improve our understanding of sensitivities to drug or genetic perturbations across cell lines.

Through our analysis, we show relevant markers of telomere-associated protein function, patterns of *TERT* reactivation across cancers, and epigenetic determinants of *TERT* promoter status. We detail various features of telomere regulation and dysfunction in cancer, and we provide a substantial addition of new features to a well-characterized set of cell lines. By doing so, we complement molecular studies of telomeres in parallel studies across the GTEx (Demanelis et al., 2019) and TCGA (Barthel et al., 2017), providing a valuable resource that will guide additional studies on the roles and functions of telomeres in cancer.

## Acknowledgements

We thank Dr. Elizabeth Blackburn (UCSF) and Dr. Thomas Cech (University of Colorado Boulder) for providing insightful comments and feedback on the manuscript. We thank Dr. Hani Goodarzi (UCSF) for his generosity in providing storage and computing resources.

## Author contributions

M.G. and F.W.H. conceived the studies. M.G. obtained initial telomere content estimates. K.H. performed all subsequent data acquisition and analysis. K.H. and F.W.H. organized figures and tables. K.H. wrote the paper, and M.G. and F.W.H. oversaw the analyses and commented on and edited the manuscript.

## Methods

### Telomere content estimation

Telomere content estimates were computed using Telseq (Ding et al., 2014) with the default settings. Telseq records the frequencies of reads containing various frequencies of the canonical TTAGGG telomeric repeat, and then normalizes this number of telomeric repeats using a GC-adjusted coverage estimate and the average chromosome length.

Telomere content was estimated for WGS and WES samples in the CCLE (Ghandi et al., 2019) as well as WES samples in the GSDC (Yang et al., 2013) using the default settings. When multiple read groups were present in a sample, telomere content was computed as a mean of the individual read group estimates weighted by the total read count per group.

Whereas we found decent agreement between overlapping samples in CCLE WGS and GDSC WES, we found a comparatively weak correlation between both sets and the CCLE WES estimates (**Supplementary Fig. 1**). Therefore, we excluded the CCLE WES telomere content estimates from subsequent analyses.

In comparing the CCLE WGS and GDSC WES estimates, we also noticed a batch effect resulting in two clusters of GDSC WES estimates. To identify and correct this batch effect across all GDSC WES estimates, we observed that these batches were distinguished by frequencies of reads containing exactly 4, 5, and 6 telomeric motifs. We then ran a k-means clustering on these read frequencies to estimate the clusters across all GDSC WES samples, which were subsequently adjusted by re-centering the mean of one cluster (after applying a z-scored log-transformation) to match the mean of the other.

We also attempted to use Telseq to estimate telomeric repeat-containing RNA (TERRA) expression across 1,019 RNA-seq samples from the CCLE. However, because the majority of these samples were found to contain little or no reads containing telomeric reads, TERRA capture was determined to be too low for any meaningful analysis.

Cell lines were annotated with sample descriptors from the CCLE data portal (Cell_lines_annotations_20181226.txt, https://portals.broadinstitute.org/ccle/data). Harmonized sample information, telomere content estimates, and other matched annotations are available in **Supplementary Table 1**.

### Genomic and transcriptomic markers

We sourced mutations and copy number estimates from the DepMap download portal (https://depmap.org/portal/download/) under the public 19Q4 release (CCLE_mutations.csv and CCLE_gene_cn.csv, respectively). We used the mutation classifications detailed in the *Variant_annotation* column.

We also downloaded processed RNAseq estimates in the form of gene expression, transcript expression, and exon inclusion estimates from the CCLE data portal under the latest release (CCLE_RNAseq_rsem_genes_tpm_20180929.txt.gz, CCLE_RNAseq_rsem_transcripts_tpm_20180929.txt.gz, and CCLE_RNAseq_ExonUsageRatio_20180929.gct.gz, respectively). We also downloaded RPPA estimates (CCLE_RPPA_20181003.csv) and global chromatin profiling results (CCLE_GlobalChromatinProfiling_20181130.csv). Before performing subsequent analyses, we transformed transcript and gene expression TPMs by taking a log_2_-transform with a pseudocount of +1. We also excluded transcripts with a standard deviation in this log_2_(TPM + 1) measure of less than 0.25 across all cell lines. We excluded exons with missing inclusion values in over 800 cell lines or with a standard deviation of less than 0.1. Pearson correlations were then used to calculate associations between gene and transcript RNA expression levels of *TERT* and *TERC* against telomere content estimates as well as other markers.

We also considered processed methylation estimates available on the CCLE data portal, namely the TSS 1kb upstream estimates as well as the promoter CpG cluster estimates, which we correlated against telomere content estimates. For these annotations, we filtered out regions with a standard deviation of less than 0.05.

Results of *TERT* and *TERC* expression associations, as well as telomere content associations, are available in **Supplementary Table 2**.

### Gene dependency associations

To identify gene dependencies associated with telomere content, we considered knockout/knockdown effects estimated in the Avana CRISPR-Cas9 (Meyers et al., 2017), Achilles RNAi (Tsherniak et al., 2017), and DRIVE RNAi (McDonald et al., 2017) datasets. For each gene dependency in each dataset, we computed the Pearson correlation coefficient against telomere content estimates generated separately with CCLE WGS and GDSC WES data. Correlation *P* values were determined using the two-tailed Student’s *t* test. All correlation coefficients and *P* values were determined using the *pearsonr* function as part of the *scipy*.*stats* Python module.

To identify codependencies with the CST complex members, we employed an iterative approach to identify highly ranked correlations. In particular, starting with a seed set of genes (the base case of which was the CST complex), we searched for codependencies between two genes *x* and *y* under the criteria that the *r*^2^ association between the two is among the top five for *x* vs. all other genes, and among the top five for *y* vs. all other genes as well. We recursively applied this method four times, which added the five shelterin components *ACD, POT1, TERF1, TERF2*, and *TINF2* to our gene set. To construct the clustered correlation matrix in **Fig. 2c**, we used the *clustermap* function as provided by the Seaborn Python library, with Ward’s method for the determination of the hierarchical clustering.

To identify significant associations between dependencies and mutations, we compared dependencies against binary categories of damaging/truncating (comprised of deleterious alterations, such as nonsense and splice-site alterations) and hotspot (highly recurrent) mutations. Using the previously-downloaded DepMap 19Q4 mutation annotations, we considered mutations as “damaging”/”truncating” if they were associated with a “damaging” label under the *Variant_annotation* column, and we considered mutations as “hotspot” if they were labeled as such in the corresponding COSMIC (Tate et al., 2019) (*isCOSMIChotspot*) or TCGA (*isTCGAhotspot*) columns. We excluded mutations with a total damaging or hotspot frequency of less than five across all profiled CCLE samples. Mutations were then compared with dependencies using a two-sided Mann-Whitney *U* test, with the two classes being non-damaging and non-hotspot mutant samples, and damaging and hotspot mutant samples, respectively.

To rank and visualize the codependencies shown in **Supplementary Fig. 6a,b** and the dependency-mutation associations shown in **Fig. 2d,e**, we used a signed *q*-value approach. We first transformed the raw false discovery rates by taking the negative of the base-10 logarithm, and we then applied a sign to this transformed value as determined by the direction of the codependency (the sign of the correlation coefficient) or dependency-mutation association (negative for greater sensitivity in mutants, and positive otherwise).

Dependency analyses results are available in **Supplementary Table 3**.

### Characterization of allele-specific TERT expression

Allele-specific expression may be detected by looking for discordant counts of reads mapping to single-nucleotide polymorphisms (SNPs) in DNA-sequencing vs. RNA-sequencing reads (Huang et al., 2015). In particular, allele-specific expression is evidenced by the biased frequency of a single allele of a heterozygous SNP in RNAseq reads compared to that of DNA-sequencing reads. To assess *TERT* expression in the context of allele-specificity, we examined cell lines for which DNA (WES or WGS) and RNA (RNAseq) sequencing data were available. To identify heterozygous anchor SNPs, we considered mutations in the *TERT* gene body called using Mutect 1.1.6 (Cibulskis et al., 2013) with default settings. We then applied a filter for mutations with at least eight reads supporting both the reference and alternate alleles that passed the Mutect quality control filter (i.e. classified as PASS). To force call the matching allele frequencies in RNA, we processed the matching aligned RNAseq reads using the ASEReadCounter tool provided in GATK 3.6 (Van der Auwera et al., 2013) with arguments - minDepth 8, --minBaseQuality 16, --minMappingQuality 255, and -U ALLOW_N_CIGAR_READS.

We then used these RNA and DNA allele frequencies to classify cell lines as monoallelic and biallelic expressors of *TERT* as well as two neighboring genes, *SLC6A19* and *CLPTM1L*. In particular, we examined the odds ratio derived from a binary contingency table with the two sets of categories being the context (DNA vs. RNA) and the allele (reference vs. alternate) of the read counts. To account for edge cases where the denominator of the odds ratio was zero, we added a pseudocount of 0.5 to each category before computing the odds ratio. We then denoted MAE lines as those having an odds ratio computed using the major allele as the denominator of greater than five. In instances where there were multiple informative SNPs, we considered only the SNP with the greatest supporting total RNA-seq read count. In cases where the same SNP was detected across multiple sources (for instance, in both CCLE WES and WGS), we considered the source with the greatest coverage of the SNP.

Allele-specific calls for *TERT, SLC6A19*, and *CLPTM1L* are described in **Supplementary Table 4**.

### Genome-wide allele-specific methylation analysis

To characterize and compare CpG-level ASM around the *TERT* genomic region, we utilized RRBS data generated by the CCLE (Ghandi et al., 2019). Mapped BAM files were downloaded from the CCLE FireCloud workspace, and ASM levels for each CpG pair were estimated using the *allelicmeth* command from the MethPipe package (Song et al., 2013). Within each sample, we first included only CpG pairs with a minimum coverage of eight reads. Next, among all 928 cell lines, CpG pairs included in less than 5% of these samples were excluded.

To estimate ASM, we employed a strategy similar to the original MethPipe ASM pipeline. For each pair of CpGs, we considered the four combinations of methylation states between the two CpGs: methylated-methylated (*mm*), methylated-unmethylated (*mu*), unmethylated-unmethylated (*uu*), and unmethylated-unmethylated (*mm*). For semi-methylated CpG pairs not subject to ASM, we would expect high and relatively equal frequencies of the *mu* and *um* pairs, whereas for ASM CpG pairs, we would expect the allele bias to result in high *mm* and *uu* counts and low *um* and *mu* counts. To quantify this imbalance, we used the mean square contingency coefficient (Φ) with a pseudocount of 0.5. Namely, for each CpG pair, we computed

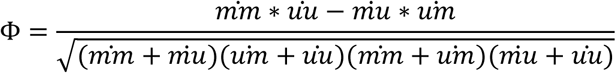

where 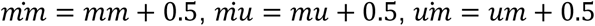, and 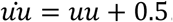. ASM CpG pairs therefore had a positive Φ, whereas non-ASM pairs had a Φ of around 0. We rounded negative Φ values to 0. Before computing these imbalance values, we excluded CpG pairs with a methylation level of less than 0.1 or greater than 0.9 on either CpG, so as to filter out CpG pairs that were likely to be fully methylated or completely demethylated.

We first examined the ASM levels of the *TERT* locus, which considered as the *TERT* gene body as well as the flanking five kilobase regions. For these methylation estimates, we excluded CpGs with less than 25% valid ASM estimates. We segmented these CpGs into five regions: the promoter (chr5:1295246-1298643), CGI_1 (chr5:1294872-1295134), CGI_2 (chr5:1291374-1294439), CGI_3 (chr5: 1289695-1291090), and the remaining gene body (chr5: 1249661-1289359). We computed ASM imbalance values for these regions by taking the mean CpG-pair ASM values.

To identify genome-wide methylation events indicative of *TERT* promoter mutations, we searched for correlates with *TERT* promoter mutation status among average methylation levels of CpG islands (CGIs). CpG island annotations were downloaded from the UCSC genome browser at http://hgdownload.cse.ucsc.edu/goldenpath/hg19/database/cpgIslandExt.txt.gz. To filter out low-coverage CpG islands, we considered only CpG islands with at least eight CpG sites. Methylation levels per island were then estimated by taking the mean across all CpGs profiled within the island. Using the same filtering parameters, we also computed mean ASM estimates across these CGIs.

*TERT* locus methylation estimates are described in **Supplementary Table 5**.

### Chromatin profiling data

To identify histone modifications associated with TERTp status, we downloaded global chromatin profiling data from the CCLE Data Portal (CCLE_GlobalChromatinProfiling_20181130.csv). Correlations between TERTp status and histone modification levels, as well as correlations between H3K9ac1K14ac0 levels and CGI ASM levels, are described in **Supplementary Table 5**.

### Region enrichment analysis

To characterize the regions that were hypomethylated in association with TERT promoter mutations, we utilized Locus Overlap Analysis (LOLA) (Nagraj et al., 2018; Sheffield & Bock, 2016), which discovers enriched regions among a background set. LOLA takes as input two region sets: the regions of interest and a background universe set. Both sets are then overlapped against a database of annotated regions, and overlapping and non-overlapping region frequencies are computed per annotated region set. Region overlap significance is then assessed against each annotated region set using Fisher’s exact test with the two categories being the regions of interest vs. the background universe set and the overlap of each of these regions with the annotated region set.

We ranked hypomethylated CpG island regions by the significance of the change in TERTp wild type vs. TERTp mutant cell lines as assessed by a two-sided Mann-Whitney *U* test (i.e. regions with the most significant changes were ranked first). The top 1,000 CpG islands were then used as the regions of interest, and the set of all CpG islands examined served as the background universe set. We utilized the LOLAweb application (at http://lolaweb.databio.org/), with the LOLACore and LOLARoadMap sets as the region databases.

Outputs of LOLA on CCLE *TERT* promoter-mutant hypomethylated CGIs are summarized in **Supplementary Table 6**.

### TCGA data

TCGA methylation and normalized gene expression estimates were downloaded from the UCSC Xena browser (Goldman et al., 2019) (http://xena.ucsc.edu/, jhu-usc.edu_PANCAN_HumanMethylation450.betaValue_whitelisted.tsv.synapse_download_50 96262.xena and EB++AdjustPANCAN_IlluminaHiSeq_RNASeqV2.geneExp.xena). Methylation levels of CGIs was estimated by averaging CpG methylation levels within each CGI, and CGIs with less than four profiled CpGs were excluded. *TERT* promoter mutation status was obtained from previous estimates (Barthel et al., 2017). The same hypomethylation-enrichment analysis previously described for the CCLE was then run on the top 1,000 CGIs hypomethylated in *TERT* promoter-mutants using an identical ranking scheme.

A summary of LOLA results on TCGA methylation, *TERT* promoter status, and ALT-likelihood is provided in **Supplementary Table 7**.

### Statistical analysis

Multiple hypothesis testing was accounted for using the Benjamini-Hochberg FDR with an alpha of 0.01 as provided by the *statsmodels* Python module.

### Data availability

Aligned CCLE sequencing data (WGS, WES, and RRBS) files were downloaded from FireCloud and are publicly available at firecloud.terra.bio/#workspaces/fccredits-silver-tan-7621/CCLE_v2, and unaligned sequencing reads were retrieved from the Sequence Read Archive (SRA) under accession PRJNA523380. GDSC WES data are available under accession number EGAD00001001039 at the European Genome-Phenome Archive (EGA). Processed CCLE and DepMap data are available at online portals as previously described.

## Supplementary figures

**Figure S1.**
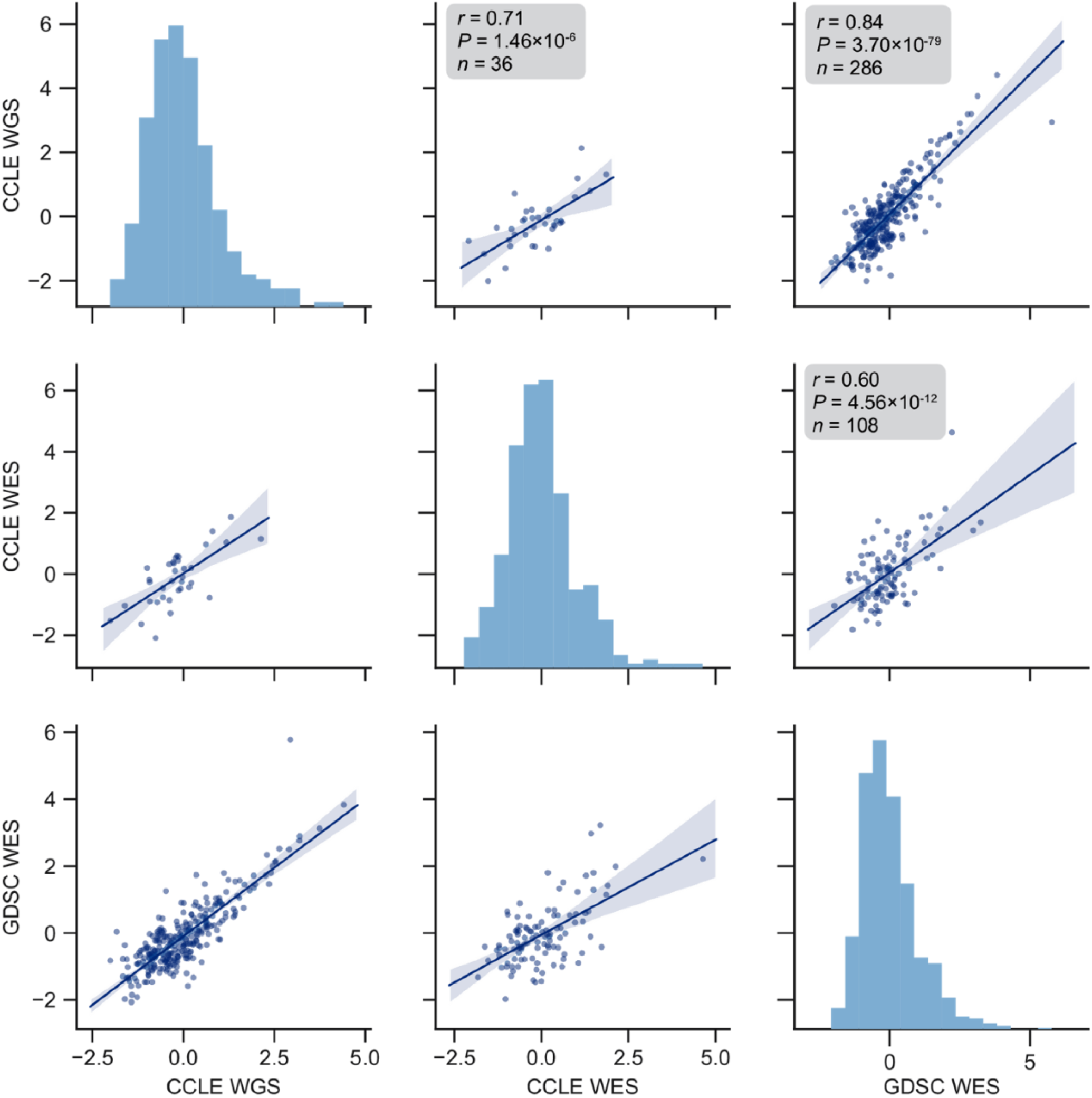
Telomere content agreement between sequencing sets. Pairwise correlations between cell lines in overlaps of the indicated sequencing sets. Telomere content estimates are displayed as *z*-scored values of log_2_-transformed raw content estimates. *P* values determined by two-sided Pearson’s correlation test.

**Figure S2.**
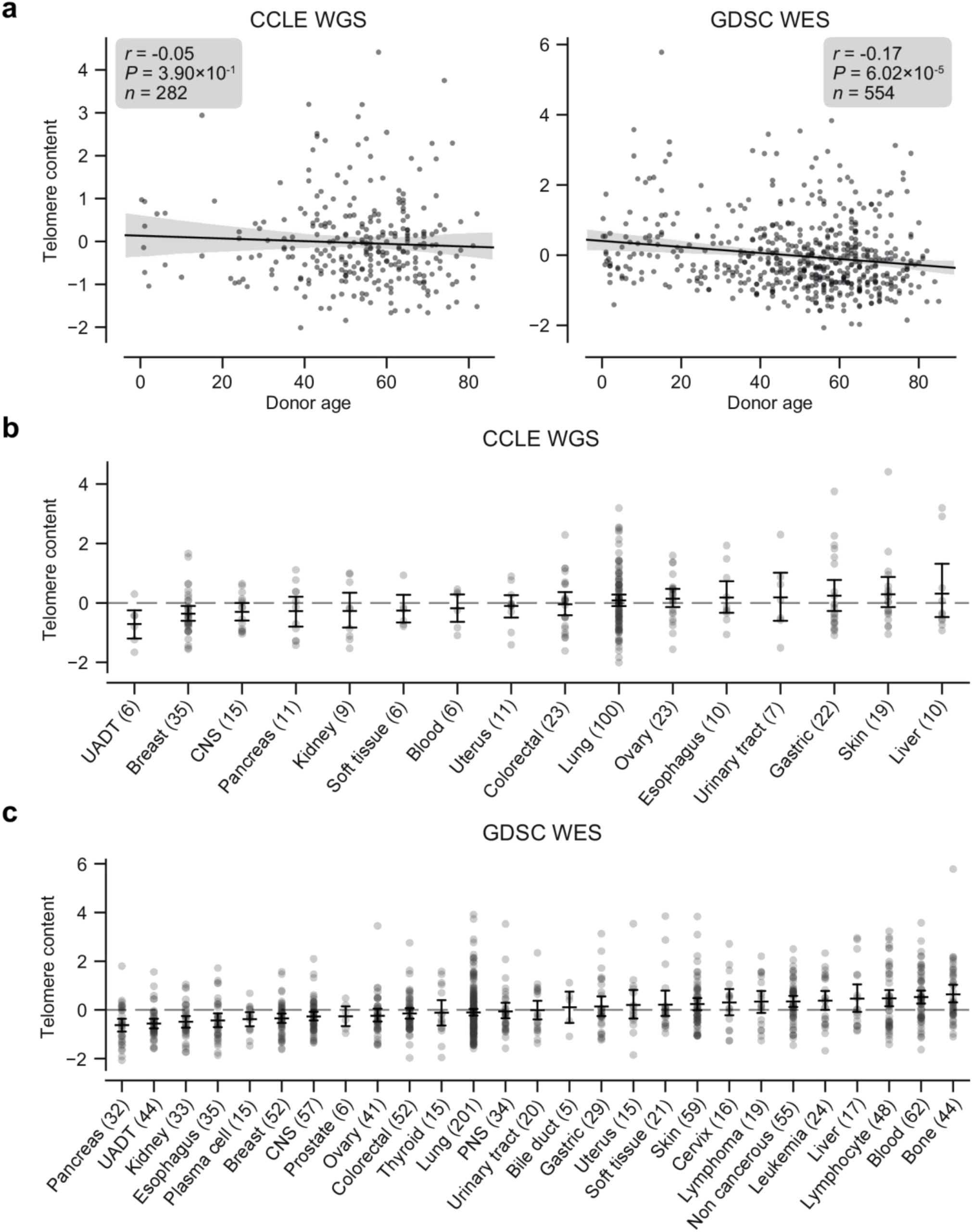
Telomere content, age, and tissue subtype. **a**, Correlation between donor age and *z*-scored log_2_ transformed telomere content estimates from CCLE WGS (left) and GDSC WES (right) telomere content estimates as labeled. **b** and **c**, Distribution of *z*-scored log_2_-transformed telomere content estimates across origin tissue subtypes of the CCLE WGS and GDSC WES datasets, respectively. Tissue subtypes with less than five samples are omitted. Bars denote bootstrapped 95% confidence intervals, with central lines denoting means. CNS, central nervous system; PNS, peripheral nervous system; UADT, upper aerodigestive tract.

**Figure S3.**
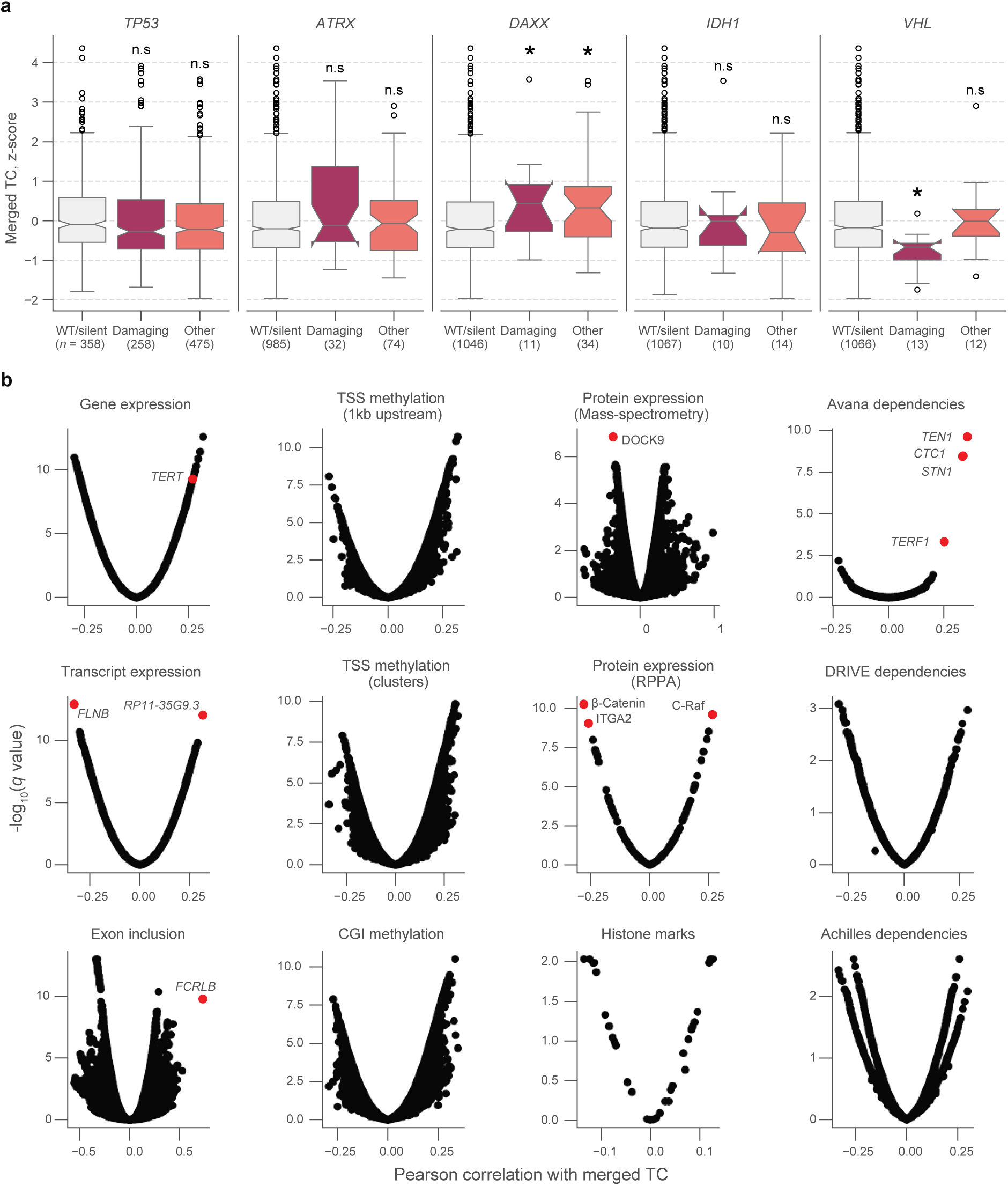
Associations between telomere content and cell line characteristics. **a**, Distributions of *z*-scored log_2_-transformed telomere content estimates from merged CCLE WGS and GDSC WES estimates, stratified by mutations in *TP53, ATRX, DAXX, IDH1*, and *VHL*. Boxes, interquartile range (IQR); center lines, median; whiskers, maximum and minimum or 1.5 × IQR; notches, 95% confidence interval of bootstrapped median using 1,000 samples and a Gaussian-based asymptotic approximation. ^*^*P* < 0.05, two-sided Mann-Whitney *U* test against WT/silent category; n.s, not significant. **b**, Volcano plots of Pearson correlations and false discovery rates (*q* values) of associations between merged telomere content estimates and several profiling datasets. Sample sizes listed in **Supplementary Table 2**. *P* values determined using two-sided Pearson’s correlation test.

**Figure S4.**
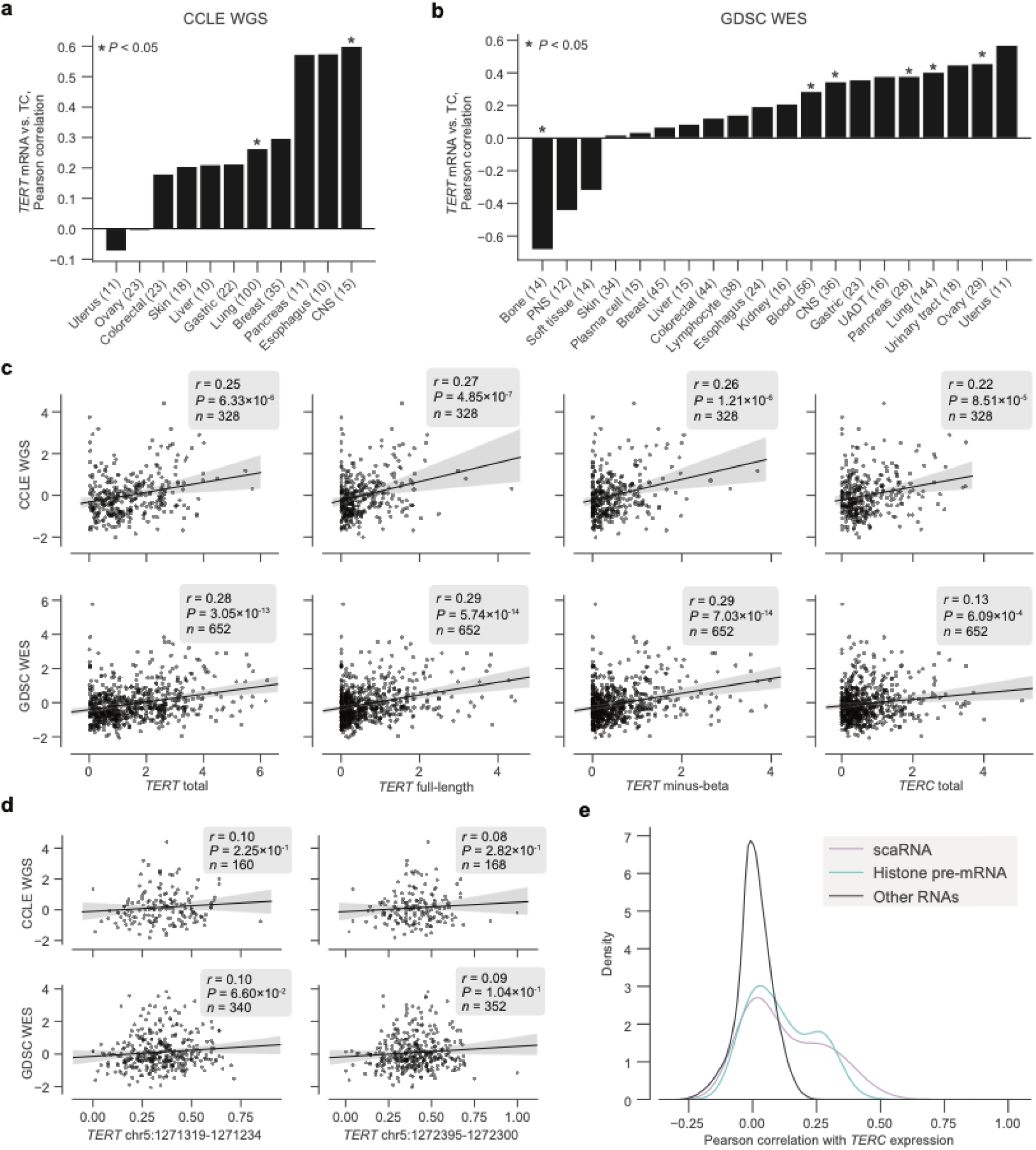
Transcriptomic associations between *TERT, TERC*, and telomere content. **a** and **b**, Pearson correlations between log_2_(TPM + 1) levels of *TERT* mRNA and telomere content within tissue subtypes in the CCLE WGS and GDSC WES datasets, respectively. CNS, central nervous system; PNS, peripheral nervous system; UADT, upper aerodigestive tract. **c**, Associations between total *TERT* (ENSG00000164362.14) mRNA, full-length *TERT* (ENST00000310581.5) mRNA, minus-beta *TERT* (ENST00000296820.5) mRNA, *TERC* (ENSG00000270141.2) RNA, and *z*-scored log_2_-transformed telomere content estimates in the CCLE WGS and GDSC WES datasets. mRNA expression measured as log_2_-transformed TPMs with a pseudocount of +1. **d**, Associations between exon inclusion levels of *TERT* exons 7 (GRCh37: chr5:1272395-1272300) and 8 (GRCh37: chr5:1271319-1271234) and telomere contents as described previously. **e**, Distribution of correlations between *TERC* RNA levels and other RNAs (*n* = 1,019 cell lines), with scaRNAs and histone pre-mRNAs highlighted.

**Figure S5.**
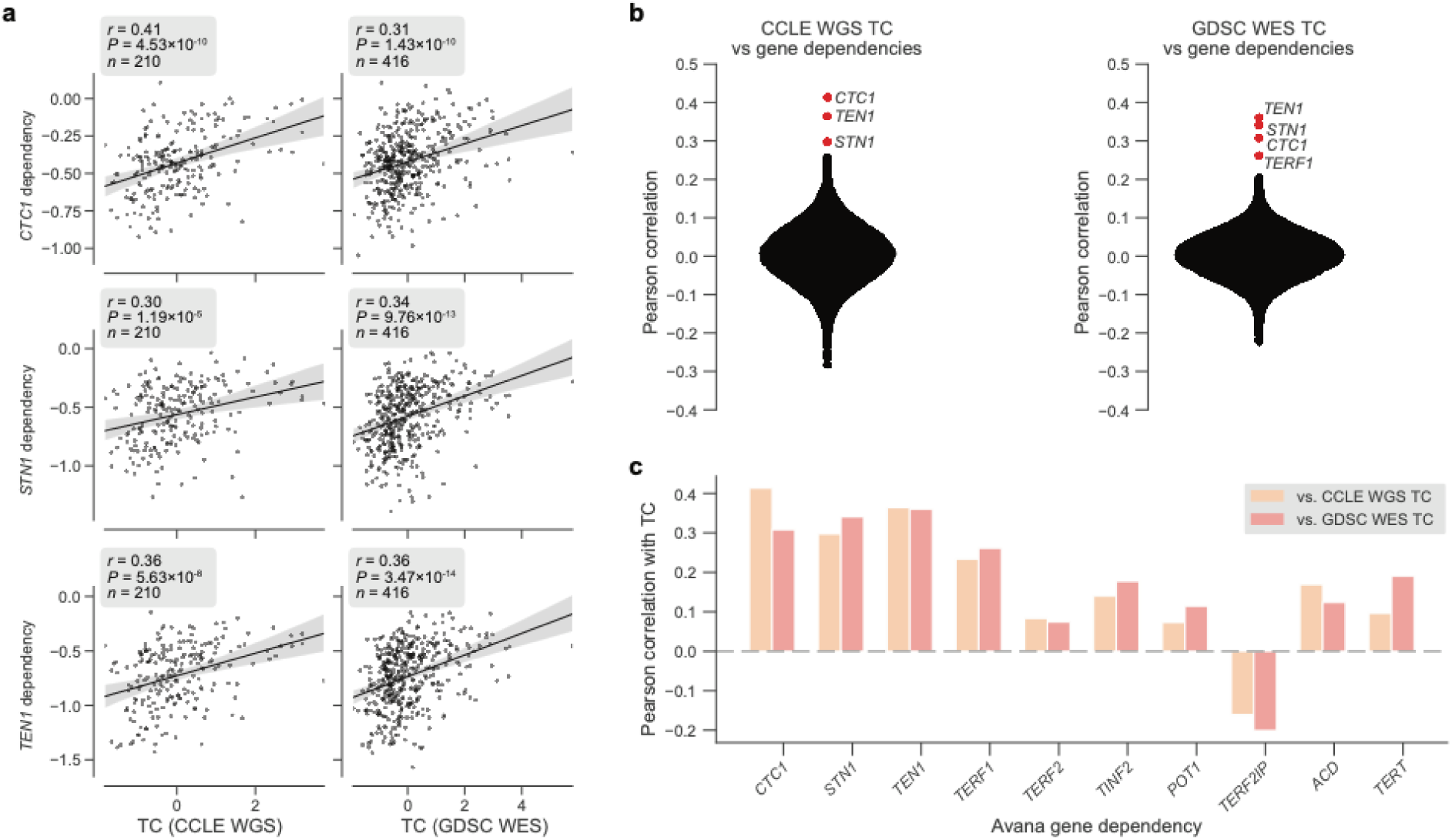
Telomere content and telomere protein dependencies. **a**, Scatterplots of *z*-scored log_2_-transformed telomere content estimates from the CCLE WGS (left) and GDSC WES (right) datasets against sensitivity to members of the CST complex measured in the Avana dataset. **b**, Distribution of Pearson correlations between telomere content estimates from CCLE WGS and GDSC WES datasets and all gene dependencies in the Avana CRISPR-Cas9 dataset (vs. CCLE WGS: *n* = 192–210 cell lines, vs. GDSC WES: *n* = 395–416 cell lines; **Supplementary Table 3**). **c**, Selected correlations between CCLE WGS and GDSC WES-derived telomere content and Avana dependencies of CST and shelterin complex members.

**Figure S6.**
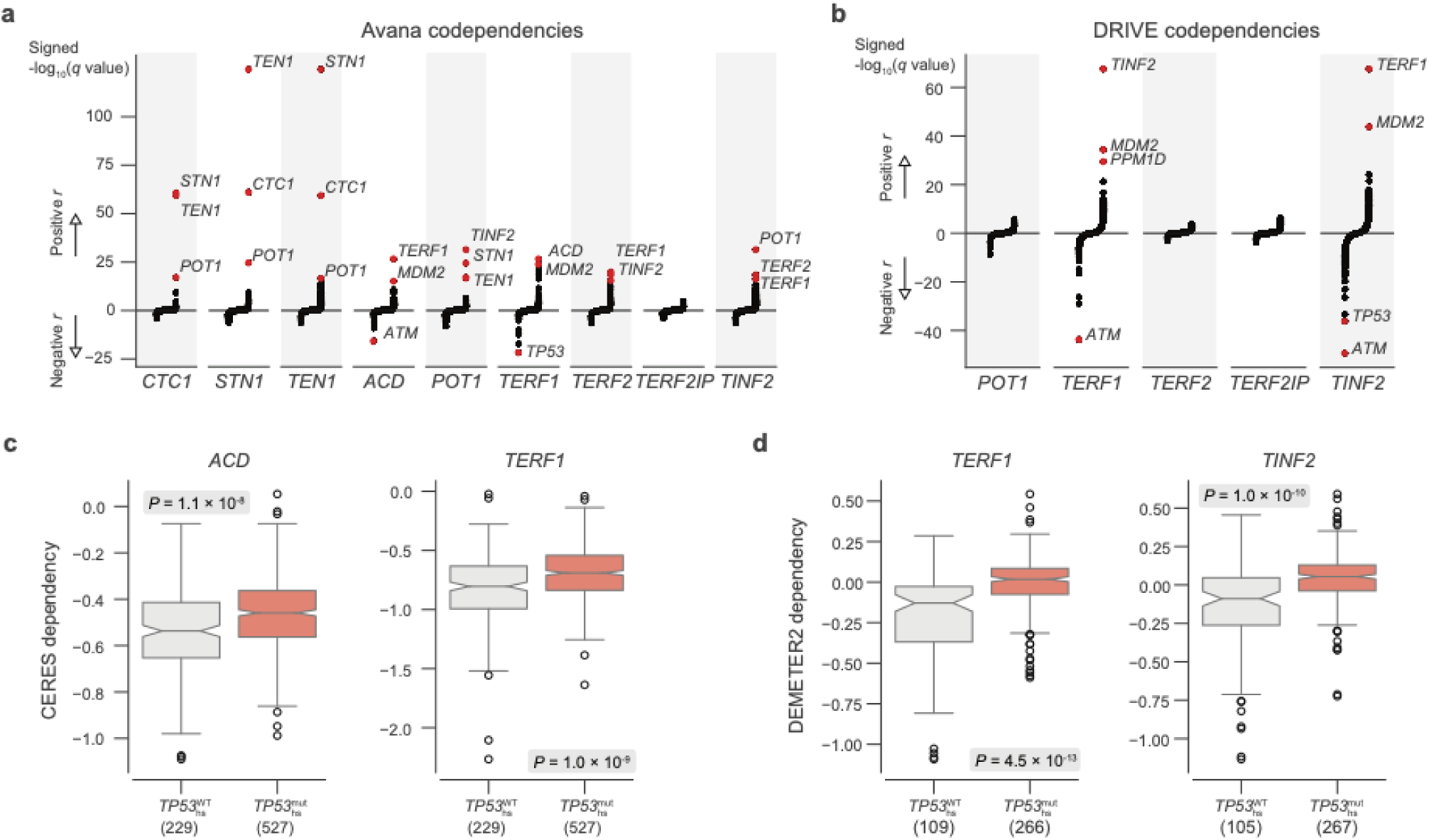
*TP53* mutation status and shelterin member dependencies. **a**, Codependencies of CST and shelterin complex members in the Avana CRISPR-Cas9 dataset as measured by Pearson correlation and the associated two-sided *P* value (*n* = 710–769 cell lines). *q*-values are shown for correlations between each gene indicated on the *x*-axis and all other genes, with *q*-values transformed and ranked by the sign of the correlation. **b**, Repeat of codependency analysis in (**a**), but for DRIVE RNAi dependencies (*n* = 88–386 cell lines). **c** and **d**, comparison of dependencies of select members (Avana and DRIVE, respectively) with respect to *TP53* hotspot mutation status. Boxes, interquartile range (IQR); center lines, median; whiskers, maximum and minimum or 1.5 × IQR; notches, 95% confidence interval of bootstrapped median using 1,000 samples and a Gaussian-based asymptotic approximation. *P* values determined by two-sided Mann-Whitney *U* test.

**Figure S7.**
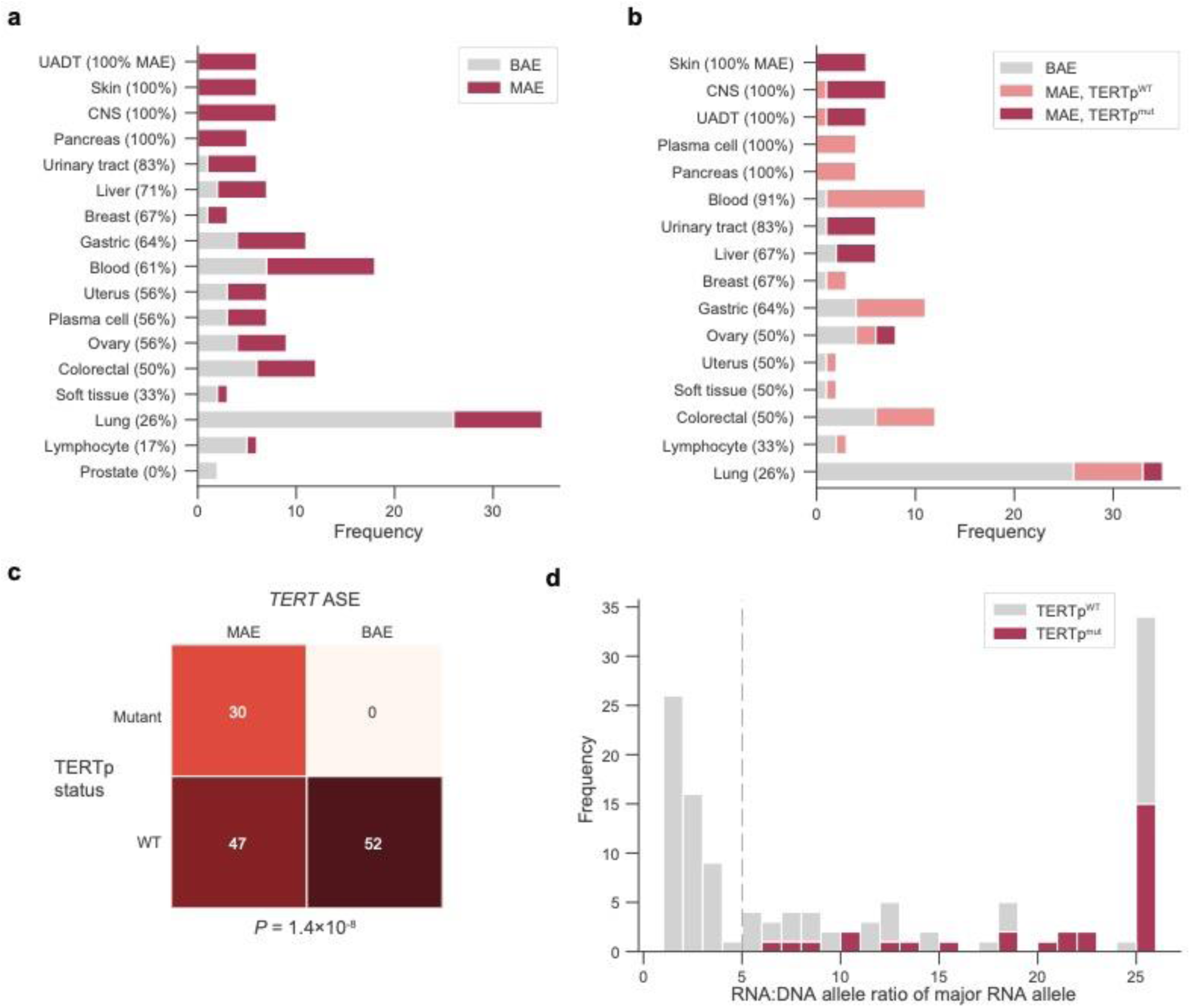
*TERT* ASE and promoter mutations. **a**, Distribution of *TERT* allele-specific expression across tissue subtypes. CNS, central nervous system; UADT, upper aerodigestive tract. **b**, Distribution of *TERT* allele-specific expression and promoter mutations across tissue subtypes. **c**, Contingency table of *TERT* allele-specific expression and *TERT* promoter mutation status. *P* value determined by Fisher’s exact test. **d**, Distribution of RNA-DNA allele ratios of the major expressed alleles in *TERT*, with matched *TERT* promoter mutation status also indicated. The dotted line at a ratio of five indicates the cutoff for classifying samples as MAE as opposed to BAE. Displayed allele ratios were capped at a maximum of 25 for viewability.

**Figure S8.**
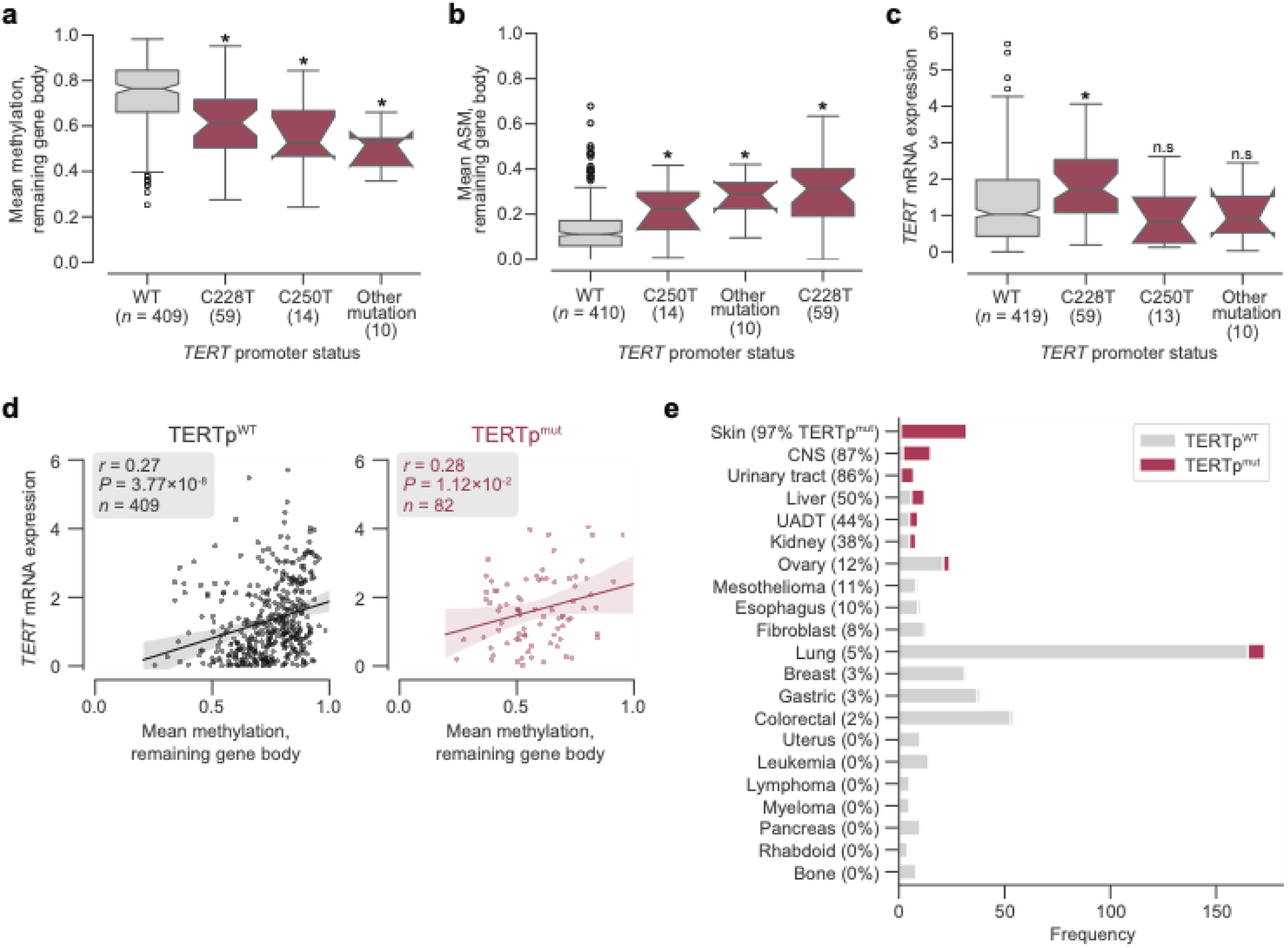
*TERT* promoter mutations, methylation, and gene expression. **a** and **b**, Distributions across different promoter mutation types of *TERT* gene body methylation (**a**), gene body ASM (**b**), and mRNA expression as measured in log_2_(TPM + 1) from RNAseq (**c**). Boxes, interquartile range (IQR); center lines, median; whiskers, maximum and minimum or 1.5 × IQR; notches, 95% confidence interval of bootstrapped median using 1,000 samples and a Gaussian-based asymptotic approximation. ^*^*P* < 0.01, two-sided Mann-Whitney *U* test against wild-type (WT) values; n.s, not significant. **d**, Correlation between *TERT* gene body methylation and mRNA expression in promoter-wildtype (left) and promoter-mutant (right) cell lines. **e**, Frequencies of *TERT* promoter mutations across different tissue subtypes.

**Figure S9.**
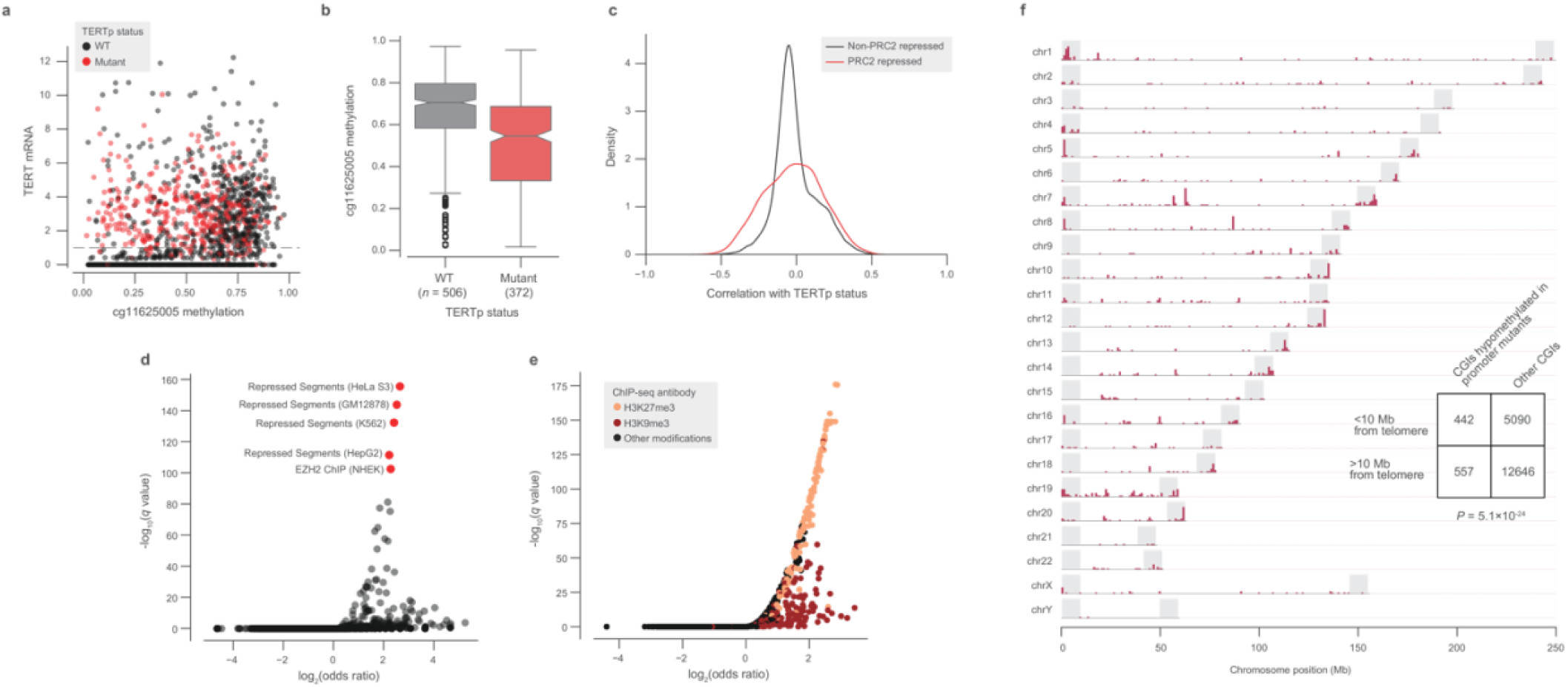
Global methylation changes associated with *TERT* promoter mutations. **a**, Methylation at the cg11625005 CpG probe and *TERT* mRNA expression across TCGA samples annotated by TERTp status (*n* = 1,553). **b**, Comparison of cg11625005 methylation levels in TERTp mutants and wild-type TCGA samples with *TERT* mRNA expression greater than 1 (as indicated in the horizontal line in (**a**). Boxes, interquartile range (IQR); center lines, median; whiskers, maximum and minimum or 1.5 × IQR; notches, 95% confidence interval of bootstrapped median using 1,000 samples and a Gaussian-based asymptotic approximation. *P* = 1.5 × 10^−26^, two-sided Mann-Whitney *U* test. **c**, Kernel density distributions of rank-biserial correlations between CGI methylation levels for PRC2-overlapping regions and non-PRC2-overlapping regions. A negative correlation indicates that a CGI is hypomethylated in TERTp^mut^ cell lines relative to TERTp^WT^ ones, and a positive correlation indicates the opposite. PRC2 regions were sourced from the HepG2 segmentation. **d**, LOLA core set enrichment analysis of CGIs hypomethylated in TERTp^mut^ samples reveals enrichment of PRC2-repressed regions. **e**, LOLA ENCODE Roadmap region enrichment analysis of CGIs hypomethylated in TERTp^mut^ samples reveals enrichment of H3K9me3 and H3K27me3 regions. **f**, Genomic distribution of CGIs hypomethylated in TERTp^mut^ CCLE cell lines. Shaded regions denote 10Mb chromosome ends.

## Supplementary tables

**Table S1**. Telomere content estimates and sample information.

**Table S2**. Telomere content and transcriptomic associations.

**Table S3**. Unsupervised dependency associations.

**Table S4**. Allele-specific expression calls.

**Table S5**. *TERT* locus methylation.

**Table S6**. Genome-wide methylation analysis in the CCLE.

**Table S7**. Genome-wide methylation analysis in TCGA.

## References

Akincilar, S. C., Khattar, E., Boon, P. L. S., Unal, B., Fullwood, M. J., & Tergaonkar, V. (2016). Long-range chromatin interactions drive mutant TERT promoter activation. Cancer Discovery, 6(11). https://doi.org/10.1158/2159-8290.CD-16-0177

Barretina, J., Caponigro, G., Stransky, N., Venkatesan, K., Margolin, A. A., Kim, S., Wilson, C. J., Lehár, J., Kryukov, G. V., Sonkin, D., Reddy, A., Liu, M., Murray, L., Berger, M. F., Monahan, J. E., Morais, P., Meltzer, J., Korejwa, A., Jané-Valbuena, J., … Garraway, L. A. (2012). The Cancer Cell Line Encyclopedia enables predictive modelling of anticancer drug sensitivity. Nature, 483(7391), 603–607. https://doi.org/10.1038/nature11003

Barthel, F. P., Wei, W., Tang, M., Martinez-Ledesma, E., Hu, X., Amin, S. B., Akdemir, K. C., Seth, S., Song, X., Wang, Q., Lichtenberg, T., Hu, J., Zhang, J., Zheng, S., & Verhaak, R. G. W. (2017). Systematic analysis of telomere length and somatic alterations in 31 cancer types. Nature Genetics, 49(3), 349–357. https://doi.org/10.1038/ng.3781

Bell, R. J. A., Rube, H. T., Kreig, A., Mancini, A., Fouse, S. D., Nagarajan, R. P., Choi, S., Hong, C., He, D., Pekmezci, M., Wiencke, J. K., Wrensch, M. R., Chang, S. M., Walsh, K. M., Myong, S., Song, J. S., & Costello, J. F. (2015). The transcription factor GABP selectively binds and activates the mutant TERT promoter in cancer. Science, 348(6238), 1036–1039. https://doi.org/10.1126/science.aab0015

Brosnan-Cashman, J. A., Yuan, M., Graham, M. K., Rizzo, A. J., Myers, K. M., Davis, C., Zhang, R., Esopi, D. M., Raabe, E. H., Eberhart, C. G., Heaphy, C. M., & Meeker, A. K. (2018). ATRX loss induces multiple hallmarks of the alternative lengthening of telomeres (ALT) phenotype in human glioma cell lines in a cell line-specific manner. PLoS ONE, 13(9). https://doi.org/10.1371/journal.pone.0204159

Bryan, T. M., Englezou, A., Dalla-Pozza, L., Dunham, M. A., & Reddel, R. R. (1997). Evidence for an alternative mechanism for maintaining telomere length in human tumors and tumor-derived cell lines. Nature Medicine, 3(11), 1271–1274. https://doi.org/10.1038/nm1197-1271

Cao, Y., Bryan, T. M., & Reddel, R. R. (2008). Increased copy number of the TERT and TERC telomerase subunit genes in cancer cells. In Cancer Science (Vol. 99, Issue 6, pp. 1092– 1099). https://doi.org/10.1111/j.1349-7006.2008.00815.x

Castel, S. E., Aguet, F., Mohammadi, P., Consortium, Gte., Ardlie, K. G., & Lappalainen, T. (2019). A vast resource of allelic expression data spanning human tissues. BioRxiv, 792911. https://doi.org/10.1101/792911

Cesare, A. J., & Reddel, R. R. (2010). Alternative lengthening of telomeres: Models, mechanisms and implications. In Nature Reviews Genetics (Vol. 11, Issue 5, pp. 319–330). https://doi.org/10.1038/nrg2763

Chen, L. Y., Redon, S., & Lingner, J. (2012). The human CST complex is a terminator of telomerase activity. Nature, 488(7412), 540–544. https://doi.org/10.1038/nature11269

Chiba, K., Johnson, J. Z., Vogan, J. M., Wagner, T., Boyle, J. M., & Hockemeyer, D. (2015). Cancer-associated tert promoter mutations abrogate telomerase silencing. ELife, 4(JULY 2015), 1–20. https://doi.org/10.7554/eLife.07918

Chiba, K., Lorbeer, F. K., Shain, A. H., McSwiggen, D. T., Schruf, E., Oh, A., Ryu, J., Darzacq, X., Bastian, B. C., & Hockemeyer, D. (2017). Mutations in the promoter of the telomerase gene TERT contribute to tumorigenesis by a two-step mechanism. Science, 357(6358), 1416–1420. https://doi.org/10.1126/science.aao0535

Cibulskis, K., Lawrence, M. S., Carter, S. L., Sivachenko, A., Jaffe, D., Sougnez, C., Gabriel, S., Meyerson, M., Lander, E. S., & Getz, G. (2013). Sensitive detection of somatic point mutations in impure and heterogeneous cancer samples. Nature Biotechnology, 31(3), 213–219. https://doi.org/10.1038/nbt.2514

Clynes, D., Jelinska, C., Xella, B., Ayyub, H., Scott, C., Mitson, M., Taylor, S., Higgs, D. R., & Gibbons, R. J. (2015). Suppression of the alternative lengthening of telomere pathway by the chromatin remodelling factor ATRX. Nature Communications, 6. https://doi.org/10.1038/ncomms8538

Damm, K., Hemmann, U., Garin-Chesa, P., Hauel, N., Kauffmann, I., Priepke, H., Niestroj, C., Daiber, C., Enenkel, B., Guilliard, B., Lauritsch, I., Müller, E., Pascolo, E., Sauter, G., Pantic, M., Martens, U. M., Wenz, C., Lingner, J., Kraut, N., … Schnapp, A. (2001). A highly selective telomerase inhibitor limiting human cancer cell proliferation. EMBO Journal, 20(24), 6958–6968. https://doi.org/10.1093/emboj/20.24.6958

De Lange, T. (2005). Shelterin: The protein complex that shapes and safeguards human telomeres. In Genes and Development (Vol. 19, Issue 18,pp. 2100–2110). https://doi.org/10.1101/gad.1346005

De Lange, T. (2009). How telomeres solve the end-protection problem. Science, 326(5955), 948–952. https://doi.org/10.1126/science.1170633

Demanelis, K., Jasmine, F., Chen, L. S., Chernoff, M., Tong, L., Shinkle, J., Sabarinathan, M., Lin, H., Ramirez, E., Oliva, M., Kim-Hellmuth, S., Stranger, B. E., Ardlie, K. G., Aguet, F., Ahsan, H., Consortium, Gte., Doherty, J., Kibriya, M. G., & Pierce, B. L. (2019). Determinants of telomere length across human tissues. BioRxiv, 793406. https://doi.org/10.1101/793406

Dikmen, Z. G., Gellert, G. C., Jackson, S., Gryaznov, S., Tressler, R., Dogan, P., Wright, W. E., & Shay, J. W. (2005). In vivo inhibition of lung cancer by GRN163L: A novel human telomerase inhibitor. Cancer Research, 65(17), 7866–7873. https://doi.org/10.1158/0008-5472.CAN-05-1215

Ding, Z., Mangino, M., Aviv, A., Spector, T., & Durbin, R. (2014). Estimating telomere length from whole genome sequence data. Nucleic Acids Research, 42(9). https://doi.org/10.1093/nar/gku181

Feng, J., Funk, W. D., Wang, S. S., Weinrich, S. L., Avilion, A. A., Chiu, C. P., Adams, R. R., Chang, E., Allsopp, R. C., Yu, J., Le, S., West, M. D., Harley, C. B., Andrews, W. H., Greider, C. W., & Villeponteau, B. (1995). The RNA component of human telomerase. Science, 269(5228), 1236–1241. https://doi.org/10.1126/science.7544491

Feuerbach, L., Sieverling, L., Deeg, K. I., Ginsbach, P., Hutter, B., Buchhalter, I., Northcott, P. A., Mughal, S. S., Chudasama, P., Glimm, H., Scholl, C., Lichter, P., Fröhling, S., Pfister, S. M., Jones, D. T. W., Rippe, K., & Brors, B. (2019). TelomereHunter - In silico estimation of telomere content and composition from cancer genomes. BMC Bioinformatics, 20(1). https://doi.org/10.1186/s12859-019-2851-0

Flynn, R. L., Cox, K. E., Jeitany, M., Wakimoto, H., Bryll, A. R., Ganem, N. J., Bersani, F., Pineda, J. R., Suvà, M. L., Benes, C. H., Haber, D. A., Boussin, F. D., & Zou, L. (2015). Alternative lengthening of telomeres renders cancer cells hypersensitive to ATR inhibitors. Science, 347(6219), 273–277. https://doi.org/10.1126/science.1257216

Gall, J. G. (2003). The centennial of the Cajal body. In Nature Reviews Molecular Cell Biology (Vol. 4, Issue 12, pp. 975–980). https://doi.org/10.1038/nrm1262

Ghandi, M., Huang, F. W., Jané-Valbuena, J., Kryukov, G. V., Lo, C. C., McDonald, E. R., Barretina, J., Gelfand, E. T., Bielski, C. M., Li, H., Hu, K., Andreev-Drakhlin, A. Y., Kim, J., Hess, J. M., Haas, B. J., Aguet, F., Weir, B. A., Rothberg, M. V., Paolella, B. R., … Sellers, W. R. (2019). Next-generation characterization of the Cancer Cell Line Encyclopedia. Nature, 569, 503–508. https://doi.org/10.1038/s41586-019-1186-3

Goldman, M., Craft, B., Hastie, M., Repečka, K., McDade, F., Kamath, A., Banerjee, A., Luo, Y., Rogers, D., Brooks, A. N., Zhu, J., & Haussler, D. (2019). The UCSC Xena platform for public and private cancer genomics data visualization and interpretation. BioRxiv, Schroeder 2015, 326470. https://doi.org/10.1101/326470

Greider, C. W. (2012). Wnt regulates TERT - Putting the horse before the cart. In Science (Vol. 336, Issue 6088, pp. 1519–1520). https://doi.org/10.1126/science.1223785

Greider, C. W., & Blackburn, E. H. (1985). Identification of a specific telomere terminal transferase activity in tetrahymena extracts. Cell, 43(2 PART 1), 405–413. https://doi.org/10.1016/0092-8674(85)90170-9

Hackett, J. A., & Greider, C. W. (2002). Balancing instability: Dual roles for telomerase and telomere dysfunction in tumorigenesis. In Oncogene (Vol. 21, Issue 4 REV. ISS. 1, pp. 619–626). https://doi.org/10.1038/sj/onc/1205061

Heaphy, C. M., De Wilde, R. F., Jiao, Y., Klein, A. P., Edil, B. H., Shi, C., Bettegowda, C., Rodriguez, F. J., Eberhart, C. G., Hebbar, S., Offerhaus, G. J., McLendon, R., Rasheed, B. A., He, Y., Yan, H., Bigner, D. D., Oba-Shinjo, S. M., Marie, S. K. N., Riggins, G. J., … Meeker, A. K. (2011). Altered telomeres in tumors with ATRX and DAXX mutations. In Science (Vol. 333, Issue 6041, p. 425). https://doi.org/10.1126/science.1207313

Heaphy, C. M., Subhawong, A. P., Hong, S. M., Goggins, M. G., Montgomery, E. A., Gabrielson, E., Netto, G. J., Epstein, J. I., Lotan, T. L., Westra, W. H., Shih, I. M., Iacobuzio-Donahue, C. A., Maitra, A., Li, Q. K., Eberhart, C. G., Taube, J. M., Rakheja, D., Kurman, R. J., Wu, T. C., … Meeker, A. K. (2011). Prevalence of the alternative lengthening of telomeres telomere maintenance mechanism in human cancer subtypes. American Journal of Pathology, 179(4), 1608–1615. https://doi.org/10.1016/j.ajpath.2011.06.018

Horn, S., Figl, A., Rachakonda, P. S., Fischer, C., Sucker, A., Gast, A., Kadel, S., Moll, I., Nagore, E., Hemminki, K., Schadendorf, D., & Kumar, R. (2013). TERT promoter mutations in familial and sporadic melanoma. Science, 339(6122), 959–961. https://doi.org/10.1126/science.1230062

Huang, F. W., Bielski, C. M., Rinne, M. L., Hahn, W. C., Sellers, W. R., Stegmeier, F., Garraway, L. A., & Kryukov, G. V. (2015). TERT promoter mutations and monoallelic activation of TERT in cancer. Oncogenesis, 4. https://doi.org/10.1038/oncsis.2015.39

Huang, F. W., Hodis, E., Xu, M. J., Kryukov, G. V., Chin, L., & Garraway, L. A. (2013). Highly recurrent TERT promoter mutations in human melanoma. Science, 339(6122), 957–959. https://doi.org/10.1126/science.1229259

Killela, P. J., Reitman, Z. J., Jiao, Y., Bettegowda, C., Agrawal, N., Diaz, L. A., Friedman, A. H., Friedman, H., Gallia, G. L., Giovanella, B. C., Grollman, A. P., He, T. C., He, Y., Hruban, R. H., Jallo, G. I., Mandahl, N., Meeker, A. K., Mertens, F., Netto, G. J., … Yan, H. (2013). TERT promoter mutations occur frequently in gliomas and a subset of tumors derived from cells with low rates of self-renewal. Proceedings of the National Academy of Sciences of the United States of America, 110(15), 6021–6026. https://doi.org/10.1073/pnas.1303607110

Kim, N. W., Piatyszek, M. A., Prowse, K. R., Harley, C. B., West, M. D., Ho, P. L. C., Coviello, G. M., Wright, W. E., Weinrich, S. L., & Shay, J. W. (1994). Specific association of human telomerase activity with immortal cells and cancer. Science, 266(5193), 2011–2015. https://doi.org/10.1126/science.7605428

Kim, W., Ludlow, A. T., Min, J., Robin, J. D., Stadler, G., Mender, I., Lai, T. P., Zhang, N., Wright, W. E., & Shay, J. W. (2016). Regulation of the Human Telomerase Gene TERT by Telomere Position Effect—Over Long Distances (TPE-OLD): Implications for Aging and Cancer. PLoS Biology, 14(12). https://doi.org/10.1371/journal.pbio.2000016

Kim, W., & Shay, J. W. (2018). Long-range telomere regulation of gene expression: Telomere looping and telomere position effect over long distances (TPE-OLD). Differentiation, 99, 1– 9. https://doi.org/10.1016/j.diff.2017.11.005

Lee, D. D., Leão, R., Komosa, M., Gallo, M., Zhang, C. H., Lipman, T., Remke, M., Heidari, A., Nunes, N. M., Apolónio, J. D., Price, A. J., De Mello, R. A., Dias, J. S., Huntsman, D., Hermanns, T., Wild, P. J., Vanner, R., Zadeh, G., Karamchandani, J., … Tabori, U. (2019). DNA hypermethylation within TERT promoter upregulates TERT expression in cancer. In Journal of Clinical Investigation (Vol. 129, Issue 4, p. 1801). American Society for Clinical Investigation. https://doi.org/10.1172/JCI121303

Lee, M., Teber, E. T., Holmes, O., Nones, K., Patch, A. M., Dagg, R. A., Lau, L. M. S., Lee, J. H., Napier, C. E., Arthur, J. W., Grimmond, S. M., Hayward, N. K., Johansson, P. A., Mann, G. J., Scolyer, R. A., Wilmott, J. S., Reddel, R. R., Pearson, J. V., Waddell, N., & Pickett, H. A. (2018). Telomere sequence content can be used to determine ALT activity in tumours. Nucleic Acids Research, 46(10), 4903–4918. https://doi.org/10.1093/nar/gky297

Leung, J. W. C., Makharashvili, N., Agarwal, P., Chiu, L. Y., Pourpre, R., Cammarata, M. B., Cannon, J. R., Sherker, A., Durocher, D., Brodbelt, J. S., Paull, T. T., & Miller, K. M. (2017). ZMYM3 regulates BRCA1 localization at damaged chromatin to promote DNA repair. Genes and Development, 31(3), 260–274. https://doi.org/10.1101/gad.292516.116

Listerman, I., Sun, J., Gazzaniga, F. S., Lukas, J. L., & Blackburn, E. H. (2013). The major reverse transcriptase-incompetent splice variant of the human telomerase protein inhibits telomerase activity but protects from apoptosis. Cancer Research, 73(9), 2817–2828. https://doi.org/10.1158/0008-5472.CAN-12-3082

Lovejoy, C. A., Li, W., Reisenweber, S., Thongthip, S., Bruno, J., de Lange, T., De, S., Petrini, J. H. J., Sung, P. A., Jasin, M., Rosenbluh, J., Zwang, Y., Weir, B. A., Hatton, C., Ivanova, E., Macconaill, L., Hanna, M., Hahn, W. C., Lue, N. F., … Meeker, A. K. (2012). Loss of ATRX, genome instability, and an altered DNA damage response are hallmarks of the alternative lengthening of Telomeres pathway. PLoS Genetics, 8(7). https://doi.org/10.1371/journal.pgen.1002772

McClintock, B. (1941). The Stability of Broken Ends of Chromosomes in Zea Mays. Genetics, 26(2), 234–282. http://www.ncbi.nlm.nih.gov/pubmed/17247004 %OA http://www.pubmedcentral.nih.gov/articlerender.fcgi?artid=PMC1209127

McClintock, B. (1942). The Fusion of Broken Ends of Chromosomes Following Nuclear Fusion. Proceedings of the National Academy of Sciences, 28(11), 458–463. https://doi.org/10.1073/pnas.28.11.458

McDonald, E. R., de Weck, A., Schlabach, M. R., Billy, E., Mavrakis, K. J., Hoffman, G. R., Belur, D., Castelletti, D., Frias, E., Gampa, K., Golji, J., Kao, I., Li, L., Megel, P., Perkins, T. A., Ramadan, N., Ruddy, D. A., Silver, S. J., Sovath, S., … Sellers, W. R. (2017). Project DRIVE: A Compendium of Cancer Dependencies and Synthetic Lethal Relationships Uncovered by Large-Scale, Deep RNAi Screening. Cell, 170(3), 577-592.e10. https://doi.org/10.1016/j.cell.2017.07.005

Meyers, R. M., Bryan, J. G., McFarland, J. M., Weir, B. A., Sizemore, A. E., Xu, H., Dharia, N. V., Montgomery, P. G., Cowley, G. S., Pantel, S., Goodale, A., Lee, Y., Ali, L. D., Jiang, G., Lubonja, R., Harrington, W. F., Strickland, M., Wu, T., Hawes, D. C., … Tsherniak, A. (2017). Computational correction of copy number effect improves specificity of CRISPR-Cas9 essentiality screens in cancer cells. Nature Genetics, 49(12), 1779–1784. https://doi.org/10.1038/ng.3984

Mitchell, J. R., Cheng, J., & Collins, K. (1999). A Box H/ACA Small Nucleolar RNA-Like Domain at the Human Telomerase RNA 3′ End. Molecular and Cellular Biology, 19(1), 567–576. https://doi.org/10.1128/mcb.19.1.567

Nagraj, V. P., Magee, N. E., & Sheffield, N. C. (2018). LOLAweb: A containerized web server for interactive genomic locus overlap enrichment analysis. Nucleic Acids Research, 46(W1), W194–W199. https://doi.org/10.1093/nar/gky464

Nusinow, D. P., Szpyt, J., Ghandi, M., Rose, C. M., McDonald, E. R., Kalocsay, M., Jané-Valbuena, J., Gelfand, E., Schweppe, D. K., Jedrychowski, M., Golji, J., Porter, D. A., Rejtar, T., Wang, Y. K., Kryukov, G. V., Stegmeier, F., Erickson, B. K., Garraway, L. A., Sellers, W. R., & Gygi, S. P. (2020). Quantitative Proteomics of the Cancer Cell Line Encyclopedia. Cell, 180(2), 387-402.e16. https://doi.org/10.1016/j.cell.2019.12.023

Olovnikov, A. M. (1973). A theory of marginotomy. The incomplete copying of template margin in enzymic synthesis of polynucleotides and biological significance of the phenomenon. Journal of Theoretical Biology, 41(1), 181–190. https://doi.org/10.1016/0022-5193(73)90198-7

Pan, J., Meyers, R. M., Michel, B. C., Mashtalir, N., Sizemore, A. E., Wells, J. N., Cassel, S. H., Vazquez, F., Weir, B. A., Hahn, W. C., Marsh, J. A., Tsherniak, A., & Kadoch, C. (2018). Interrogation of Mammalian Protein Complex Structure, Function, and Membership Using Genome-Scale Fitness Screens. Cell Systems, 6(5), 555-568.e7. https://doi.org/10.1016/j.cels.2018.04.011

Pereboeva, L., Hubbard, M., Goldman, F. D., & Westin, E. R. (2016). Robust DNA damage response and elevated reactive oxygen species in TINF2-mutated dyskeratosis congenita cells. PLoS ONE, 11(2). https://doi.org/10.1371/journal.pone.0148793

Ramamoorthy, M., & Smith, S. (2015). Loss of ATRX Suppresses Resolution of Telomere Cohesion to Control Recombination in ALT Cancer Cells. Cancer Cell, 28(3), 357–369. https://doi.org/10.1016/j.ccell.2015.08.003

Rosenfeld, J. A., Wang, Z., Schones, D. E., Zhao, K., DeSalle, R., & Zhang, M. Q. (2009). Determination of enriched histone modifications in non-genic portions of the human genome. BMC Genomics, 10. https://doi.org/10.1186/1471-2164-10-143

Rowland, T. J., Bonham, A. J., & Cech, T. R. (2020). Allele-specific proximal promoter hypomethylation of the telomerase reverse transcriptase gene (TERT) associates with TERT expression in multiple cancers. Molecular Oncology, 14(10). https://doi.org/10.1002/1878-0261.12786

Rowland, T. J., Dumbović, G., Hass, E. P., Rinn, J. L., & Cech, T. R. (2019). Single-cell imaging reveals unexpected heterogeneity of telomerase reverse transcriptase expression across human cancer cell lines. Proceedings of the National Academy of Sciences of the United States of America, 116(37), 18488–18497. https://doi.org/10.1073/pnas.1908275116

Ruthenburg, A. J., Li, H., Patel, D. J., & David Allis, C. (2007). Multivalent engagement of chromatin modifications by linked binding modules. In Nature Reviews Molecular Cell Biology (Vol. 8, Issue 12, pp. 983–994). https://doi.org/10.1038/nrm2298

Salgado, C., Roelse, C., Nell, R., Gruis, N., Doorn R. van, & Velden P. van der. (2019). Interplay between TERT promoter mutations and methylation culminates in chromatin accessibility and TERT expression. BioRxiv, 859892. https://doi.org/10.1101/859892

Scheel, C., Schaefer, K. L., Jauch, A., Keller, M., Wai, D., Brinkschmidt, C., Van Valen, F., Boecker, W., Dockhorn-Dworniczak, B., & Poremba, C. (2001). Alternative lengthening of telomeres is associated with chromosomal instability in osteosarcomas. Oncogene, 20(29), 3835–3844. https://doi.org/10.1038/sj.onc.1204493

Sheffield, N. C., & Bock, C. (2016). LOLA: Enrichment analysis for genomic region sets and regulatory elements in R and Bioconductor. Bioinformatics, 32(4), 587–589. https://doi.org/10.1093/bioinformatics/btv612

Song, Q., Decato, B., Hong, E. E., Zhou, M., Fang, F., Qu, J., Garvin, T., Kessler, M., Zhou, J., & Smith, A. D. (2013). A reference methylome database and analysis pipeline to facilitate integrative and comparative epigenomics. PLoS ONE, 8(12). https://doi.org/10.1371/journal.pone.0081148

Stern, J. L., Paucek, R. D., Huang, F. W., Ghandi, M., Nwumeh, R., Costello, J. C., & Cech, T. R. (2017). Allele-Specific DNA Methylation and Its Interplay with Repressive Histone Marks at Promoter-Mutant TERT Genes. Cell Reports, 21(13), 3700–3707. https://doi.org/10.1016/j.celrep.2017.12.001

Stern, J. L., Theodorescu, D., Vogelstein, B., Papadopoulos, N., & Cech, T. R. (2015). Mutation of the TERT promoter, switch to active chromatin, and monoallelic TERT expression in multiple cancers. Genes and Development, 29(21), 2219–2224. https://doi.org/10.1101/gad.269498.115

Takakura, M., Kyo, S., Kanaya, T., Tanaka, M., & Inoue, M. (1998). Expression of human telomerase subunits and correlation with telomerase activity in cervical cancer. Cancer Research, 58(7), 1558–1561.

Tao, Y., Kang, B., Petkovich, D. A., Bhandari, Y. R., In, J., Stein-O’Brien, G., Kong, X., Xie, W., Zachos, N., Maegawa, S., Vaidya, H., Brown, S., Chiu Yen, R. W., Shao, X., Thakor, J., Lu, Z., Cai, Y., Zhang, Y., Mallona, I., … Easwaran, H. (2019). Aging-like Spontaneous Epigenetic Silencing Facilitates Wnt Activation, Stemness, and Braf V600E -Induced Tumorigenesis. Cancer Cell, 35(2), 315-328.e6. https://doi.org/10.1016/j.ccell.2019.01.005

Tate, J. G., Bamford, S., Jubb, H. C., Sondka, Z., Beare, D. M., Bindal, N., Boutselakis, H., Cole, C. G., Creatore, C., Dawson, E., Fish, P., Harsha, B., Hathaway, C., Jupe, S. C., Kok, C. Y., Noble, K., Ponting, L., Ramshaw, C. C., Rye, C. E., … Forbes, S. A. (2019). COSMIC: The Catalogue Of Somatic Mutations In Cancer. Nucleic Acids Research, 47(D1), D941–D947. https://doi.org/10.1093/nar/gky1015

Tsherniak, A., Vazquez, F., Montgomery, P. G., Weir, B. A., Kryukov, G., Cowley, G. S., Gill, S., Harrington, W. F., Pantel, S., Krill-Burger, J. M., Meyers, R. M., Ali, L., Goodale, A., Lee, Y., Jiang, G., Hsiao, J., Gerath, W. F. J., Howell, S., Merkel, E., … Hahn, W. C. (2017). Defining a Cancer Dependency Map. Cell, 170(3), 564-576.e16. https://doi.org/10.1016/j.cell.2017.06.010

Van der Auwera, G. A., Carneiro, M. O., Hartl, C., Poplin, R., del Angel, G., Levy-Moonshine, A., Jordan, T., Shakir, K., Roazen, D., Thibault, J., Banks, E., Garimella, K. V., Altshuler, D., Gabriel, S., & DePristo, M. A. (2013). From fastQ data to high-confidence variant calls: The genome analysis toolkit best practices pipeline. Current Protocols in Bioinformatics, 43(1), 11.10.1-11.10.33. https://doi.org/10.1002/0471250953.bi1110s43

Venteicher, A. S., Abreu, E. B., Meng, Z., McCann, K. E., Terns, R. M., Veenstra, T. D., Terns, M. P., & Artandi, S. E. (2009). A human telomerase holoenzyme protein required for Cajal body localization and telomere synthesis. Science, 323(5914), 644–648. https://doi.org/10.1126/science.1165357

Verdun, R. E., & Karlseder, J. (2007). Replication and protection of telomeres. In Nature (Vol. 447, Issue 7147, pp. 924–931). Nature Publishing Group. https://doi.org/10.1038/nature05976

Wan, M., Qin, J., Songyang, Z., & Liu, D. (2009). OB fold-containing protein 1 (OBFC1), a human homolog of yeast Stn1, associates with TPP1 and is implicated in telomere length regulation. Journal of Biological Chemistry, 284(39), 26725–26731. https://doi.org/10.1074/jbc.M109.021105

Wu, K. J., Grandori, C., Amacker, M., Simon-Vermot, N., Polack, A., Lingner, J., & Dalla-Favera, R. (1999). Direct activation of TERT transcription by c-MYC. Nature Genetics, 21(2), 220– 224. https://doi.org/10.1038/6010

Yang, W., Soares, J., Greninger, P., Edelman, E. J., Lightfoot, H., Forbes, S., Bindal, N., Beare, D., Smith, J. A., Thompson, I. R., Ramaswamy, S., Futreal, P. A., Haber, D. A., Stratton, M. R., Benes, C., McDermott, U., & Garnett, M. J. (2013). Genomics of Drug Sensitivity in Cancer (GDSC): A resource for therapeutic biomarker discovery in cancer cells. Nucleic Acids Research, 41(D1). https://doi.org/10.1093/nar/gks1111

Yu, G. L., Bradley, J. D., Attardi, L. D., & Blackburn, E. H. (1990). In vivo alteration of telomere sequences and senescence caused by mutated Tetrahymena telomerase RNAs. Nature, 344(6262), 126–132. https://doi.org/10.1038/344126a0

Yuan, X., Larsson, C., & Xu, D. (2019). Mechanisms underlying the activation of TERT transcription and telomerase activity in human cancer: old actors and new players. In Oncogene (Vol. 38, Issue 34, pp. 6172–6183). https://doi.org/10.1038/s41388-019-0872-9

Zhang, A., Zheng, C., Lindvall, C., Hou, M., Ekedahl, J., Lewensohn, R., Yan, Z., Yang, X., Henriksson, M., Blennow, E., Nordenskjold, M., Zetterberg, A., Bjorkholm, M., Gruber, A., & Xu, D. (2000). Frequent amplification of the telomerase reverse transcriptase gene in human tumors. Cancer Research, 60(22), 6230–6235.

Zhu, Y., Tomlinson, R. L., Lukowiak, A. A., Terns, R. M., & Terns, M. P. (2004). Telomerase RNA Accumulates in Cajal Bodies in Human Cancer Cells. Molecular Biology of the Cell, 15(1), 81–90. https://doi.org/10.1091/mbc.E03-07-0525

